# Coregulated assembly of actin-like FtsA polymers with FtsZ during Z-ring formation and division in *Escherichia coli*

**DOI:** 10.1101/2021.01.12.426377

**Authors:** Josiah J. Morrison, Joseph Conti, Jodi L. Camberg

## Abstract

In *Escherichia coli*, the actin homolog FtsA localizes the cell division machinery, beginning with the Z-ring, to the cytoplasmic membrane through direct interaction with FtsZ. FtsZ polymers are first to assemble at the Z-ring at midcell, where they direct constriction and septation. While FtsZ polymerization is critical for establishing a functional Z-ring that leads to constriction, the assembly state of FtsA and the role of FtsA ATP utilization during division in *E. coli* remain unclear. Here, we show that ATP hydrolysis, FtsZ interaction, and phospholipid vesicle remodeling by FtsA are impaired by a substitution mutation at the predicted active site for hydrolysis. This mutation, Glu 14 to Arg, also impairs Z-ring assembly and division in vivo. To further investigate the role of phospholipid engagement and ATP utilization in regulating FtsA function, we characterized a truncated *E. coli* FtsA variant, FtsA(ΔMTS), which lacks the region at the C-terminus important for engaging the membrane and is defective for ATP hydrolysis. We show that *E. coli* FtsA(ΔMTS) forms ATP-dependent actin-like filaments and assembly is antagonized by FtsZ. Polymerization of FtsZ with GTP, or a non-hydrolyzable analog, blocks inhibition of ATP-dependent FtsA assembly, and instead favors coassembly of stable FtsA/FtsZ polymers. In the cell, FtsA/FtsZ coassembly is favored at midcell, where FtsZ polymerizes, and inhibited at regions where FtsZ polymers are destabilized by regulators, such as MinC at the poles or SlmA at the nucleoid. We show that MinC prevents recruitment of FtsZ, via FtsA, to phospholipids, suggesting that local interactions of MinC with FtsZ block membrane tethering and uncouple the Z-ring from its major membrane contact. During Z-ring formation, the coassembly of FtsZ polymers with FtsA is coordinated and is a critical early step in division. This step also serves as a checkpoint by responding to the suite of FtsZ assembly regulators in the cell that modulate Z-ring position and dynamics prior to initiating cell wall synthesis.

## Introduction

Cell division in bacteria occurs through the organized and concerted efforts of many proteins that temporally localize to the division site and execute a physical process to divide a single cell into two discrete cells. The architectural remodeling that occurs at the site of septation during division in *Escherichia coli* initiates with the assembly of the Z-ring at midcell. The most conserved and essential protein of the divisome, FtsZ, forms highly dynamic polymers that locate to the center of the cell and may serve as a scaffold to facilitate cell constriction and septum formation [1]. FtsZ polymers are unable to bind to the inner membrane of the cell directly, and two proteins, FtsA and the less conserved ZipA, serve as membrane tethers for FtsZ [2-7]. FtsA is a highly conserved actin homolog that interacts directly with the FtsZ C-terminus [7, 8]. FtsA binds to the inner membrane through a conserved C-terminal amphipathic helix that is required for in vivo function [4]. While FtsA hydrolyzes ATP rapidly in vitro in a reaction that requires phospholipids [7], the cellular role or requirement for ATP has not been clearly elucidated. In addition to its ATP hydrolyzing activity, FtsA remodels and tubulates liposomes, destabilizes FtsZ polymers in vitro, and modifies FtsZ dynamics in a reconstituted system [7, 9]. FtsA and FtsZ act as the first of at least 12 proteins that localize to midcell during the early steps of division in *E. coli* [10-13]. FtsA interacts with many other cell division proteins directly including ZipA, FtsI, FtsEX, and, most notably, FtsN [14-17].The interaction between FtsA and FtsN has been suggested to serve as a trigger to engage FtsQBL, which activates FtsWI complex for peptidoglycan (PG) synthesis [18].

In eukaryotes, actin polymerizes intracellularly to provide a networked cytoskeletal structure, aids in protein locomotion, trafficking, and cell division, and is the basis for muscle contraction in higher eukaryotes [19-22]. Notable bacterial actin homologs include the plasmid segregation protein ParM, cell shape determining protein MreB, and cell division protein FtsA [23-25]. FtsA contains similar domain organization to actin; however, the 1C domain of FtsA is in a different orientation as the 1B domain of actin and participates in direct interactions with later stage cell division proteins [8, 14-17, 24]. This domain change has significant implications for the structure of putative FtsA filaments. Indeed, crystallization of an FtsA dimer from *Thermotoga maritima* shows that dimerization appears to occur through the 1A and 2B domains near the active site interacting with the 1C and 2A domains on a subsequent protomer, whereas MreB self-interacts similarly to actin through repeated binding of domains 2B and 1B with 2A [8, 24]. Actin polymerization is dependent on ATP but is also modulated by ionic strength and divalent cations; in addition, actin assembly is regulated by a suite of other eukaryotic proteins [26-29]. Recently, high resolution cryo-EM conformational studies have revealed major structural changes for ATP-bound actin in a polymer, while minor, but notable, changes take place upon ATP hydrolysis and phosphate release [30, 31]. Although actin filament formation has been well studied, direct confirmation of *E. coli* FtsA polymerization has been elusive. After early reports of *E. coli* FtsA self-interaction from yeast two-hybrid studies [32], Pichoff and Lutkenhaus reported that a Gfp-FtsA fusion protein lacking the C-terminal membrane targeting sequence, Gfp-FtsA(Δ15), formed fluorescent rod-like structures in the cytoplasm [4], and mutants defective for rod formation were also defective for self-interaction in a yeast two-hybrid system [33]. Interestingly, cells expressing mutant FtsA proteins defective for self-interaction bypass the requirement for ZipA, as well as FtsEX, but are dependent on FtsN, suggesting that these proteins may help promote monomerization of FtsA in vivo [17, 34-37]. In vitro FtsA protofilament-like structures have been observed by transmission electron microscopy (TEM) on lipid monolayers of FtsA from *T. maritima* [8] and *E. coli* [38, 39] and in solution of FtsA from *S. pneumoniae* [40] and *V. cholerae* [41]. Only *S. pneumoniae* FtsA has shown ATP-dependent polymerization [40]. Moreover, mechanistic detail into FtsA polymerization, depolymerization and the ATP cycle remains undetermined.

Several models have been proposed to resolve the precise individual roles for FtsZ and FtsA during cell division, including but not limited to scaffolds, constriction machines, and spatiotemporal organizers. While there is a large body of evidence to suggest that FtsZ assembles into tubulin-like protofilaments in vivo, there is scant evidence to demonstrate that *E. coli* FtsA also polymerizes as part of its functional role. Furthermore, both FtsZ and FtsA localize to a nascent Z-ring in vivo, thus creating a paradox about the initial step of divisome assembly [42]. Here, we demonstrate that *E. coli* FtsA lacking the C-terminal membrane targeting sequence (MTS), FtsA(ΔMTS), forms ATP-dependent, linear polymers. Furthermore, substitution of a critical residue in the FtsA active site, E14, leads to impaired ATP hydrolysis, reduced FtsZ interactions in vitro and significant functional defects in vivo. Our results show that FtsA polymerization is coregulated with FtsZ polymerization, and that the FtsA-FtsZ interaction is disrupted by MinC, which destabilizes FtsZ polymers. Consistent with this, deletion of *minC* in vivo leads to mislocalized Z-rings in vivo, and also mislocalized Gfp-FtsA, indicating that the spatiotemporal regulation of FtsZ polymerization is conveyed to FtsA. In our model, the FtsZ polymerization state is coordinated with FtsA, and the FtsA/FtsZ coassembled state is then communicated to inner membrane protein FtsN. We predict that other cell division proteins, such as ZipA or FtsEX may contribute to regulating the FtsA polymerization state [17, 43-45]. Once recruited by FtsA, FtsN would then activate FtsQBL and FtsWI for cell wall synthesis [18, 45].

## Results

### FtsA(E14R) is defective for phospholipid remodeling and FtsZ interactions

In *E. coli* FtsA, which is modeled on the crystal structure of *T. maritima* FtsA (pdb: 4A2B) [8], Glu 14 contacts Mg^2+^ in the active site (Fig. 1A). Therefore, we predicted that E14 would be important for FtsA function, possibly through nucleotide sensing or hydrolysis activity. In an alignment of *E. coli* FtsA with hexokinase, Hsc70-like proteins, and several actins, this residue is highly conserved as a negatively charged amino acid [46]. To determine if Glu 14 is important for FtsA function in *E. coli*, we first tested if an FtsA substitution mutant, constructed as a Gfp fusion protein, localizes to Z-rings in vivo. We mutagenized Glu14 to Arg in a plasmid that encodes a Gfp-FtsA fusion protein (pSEB293) [4] and expressed the wild type and mutant fusion proteins in *E. coli* MG1655 (Fig. 1B and S1A). As expected, Gfp-FtsA exhibited a fluorescence localization pattern consistent with a division ring at midcell (Fig. 1B). Cells with the plasmid encoding Gfp-FtsA(E14R) contained solely cytoplasmic fluorescence, and we detected no ring or foci formation in these cells (Fig. 1B). FtsA contains a membrane targeting sequence (MTS) at its C-terminus that localizes FtsA to the cytoplasmic membrane at the divisome in vivo [4, 47] and directly recruits FtsA to phospholipids in vitro [7]. The MTS penetrates into the nonpolar core of the lipid bilayer, and this insertion is required for rapid FtsA ATP hydrolysis in vitro [7]. Deletion of the MTS from FtsA or removal of phospholipids from reactions containing wild type FtsA abrogates rapid ATPase activity suggesting that membrane association is critical for FtsA enzyme function and regulation [7]. We constructed and expressed Gfp-FtsA(ΔMTS) and confirmed that it was unable to localize to a ring but instead formed long, narrow foci adjacent to the cell membrane, which have been previously reported [4, 44, 48]. The E14R mutation was introduced into the plasmid encoding Gfp-FtsA(ΔMTS) to construct Gfp-FtsA(E14R/ΔMTS). In contrast to the long, narrow foci exhibited by Gfp-FtsA(ΔMTS), we instead observed diffuse cytoplasmic fluorescence (Fig. 1B). The fluorescent rod-like structures of Gfp-FtsA(ΔMTS) observed in vivo were previously hypothesized to report FtsA(ΔMTS) self-interaction, and this phenotype has been used to identify self-interaction deficient FtsA mutants in vivo [4, 33]. Our results suggest that introduction of the E14R mutation therefore prevents self-association of Gfp-FtsA(ΔMTS) (Fig. 1B).

**Fig 1.**
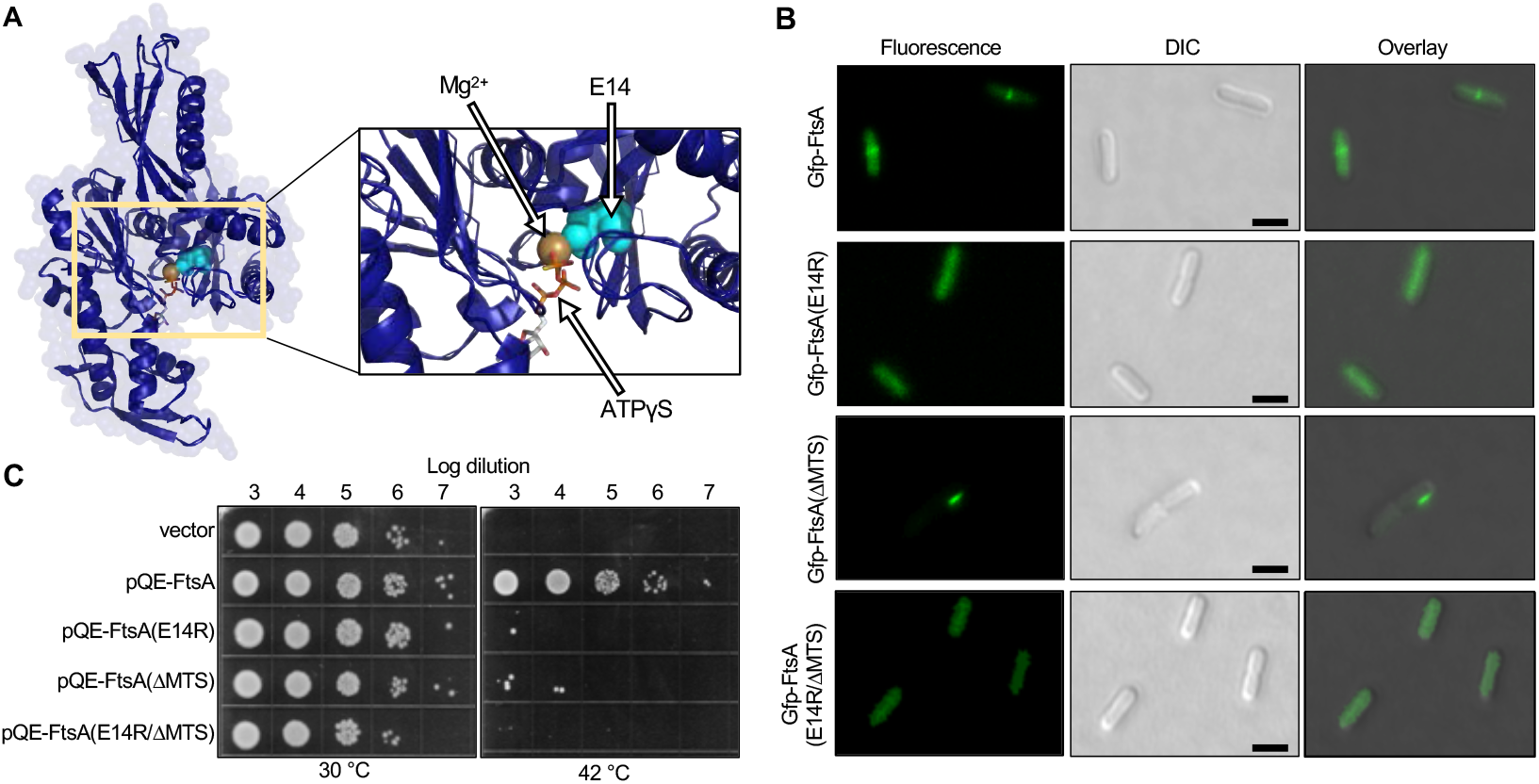
FtsA(E14R), which contains a mutation close to the active site, is defective for function in vivo. (A) *E. coli* FtsA (residues 13-381) modeled onto *T. maritima* FtsA (pdb: 4A2B) [8]. Amino acid Glu 14 (cyan) and Mg^2+^ divalent cation (gold) are shown as Corey-Pauling-Koltun (CPK) models. ATPγS is shown as stick model. (B) Confocal fluorescence and differential interference contrast (DIC) microscopy of *E. coli* MG1655 cells expressing plasmid-encoded Gfp-FtsA, Gfp-FtsA(E14R), Gfp-FtsA(ΔMTS), or Gfp-FtsA(E14R/ΔMTS). Cells were grown as described in Materials and Methods. Scale bars are 2 µm. (C) Log dilutions of cultured *E. coli* MCA12 (*ftsA*12) cells expressing FtsA or FtsA mutant proteins [FtsA(E14R), FtsA(ΔMTS), FtsA(E14R/ΔMTS) from a plasmid and grown overnight at permissive (30 °C) and restrictive (42 °C) temperatures, where indicated, on LB agar plates containing ampicillin (100 μg ml^−1^). Data shown is representative of three replicates.

Next, to determine if FtsA(E14R) supports division in vivo, we expressed the mutant protein in two temperature-sensitive strains, MCA12 and MCA27 [49], and tested if plasmid-encoded FtsA(E14R) restores growth at the restrictive temperature. MCA12 contains a chromosomal *ftsA* mutation that substitutes amino acid Ala 188 with Val, and MCA27 contains a chromosomal *ftsA* mutation that substitutes amino acid Ser 195 with Pro [49]. Cultures of temperature-sensitive strains containing plasmids encoding FtsA and FtsA(E14R) were grown at 30 °C to log phase (OD_600_ 0.4 A.U.), diluted and spotted onto LB agar plates containing ampicillin and grown overnight at the permissive (30 °C) and restrictive (42 °C) temperatures. We observed that the strains containing the vector (pQE-9) were unable to grow at the restrictive temperature (Fig. 1C and Fig. S1B). However, growth at high temperature was restored in both strains by the introduction of a plasmid containing *ftsA* (Fig. 1C and Fig.S1B). A plasmid encoding FtsA(E14R) did not support growth of either strain at the restrictive temperature, although all strains grew at the permissive temperature (Fig. 1C and Fig.S1B). A plasmid encoding FtsA(ΔMTS) also did not support growth at the restrictive temperature, which is in agreement with a previous report [4]. Finally, a plasmid containing both mutations in *ftsA*, encoding the E14R substitution and the premature stop codon to remove the MTS, was unable to restore growth of either MCA12 or MCA27 at the restrictive temperature (Fig. 1C and Fig.S1B). These results suggest that FtsA(E14R), FtsA(ΔMTS) and FtsA(E14R/ΔMTS) are non-functional in vivo. Together, our results indicate that not only does FtsA(E14R) show no localization to the Z-ring and disrupted rod formation as a fluorescent fusion protein, an expression plasmid encoding FtsA(E14R) it is also unable to complement temperature sensitive strains in vivo suggesting that this protein has major functional defects.

To investigate why FtsA(E14R) is defective for function and localization in vivo, we purified FtsA(E14R) to determine if it is also defective for known FtsA activities in vitro. The failure to assemble into foci-like structures adjacent to the cell membrane, or at a division ring, could suggest that the protein is impaired for self-interaction, phospholipid binding, or ATP hydrolysis. To determine if FtsA(E14R) is capable of robust ATP hydrolysis similar to wild type FtsA, we measured the ATPase activity and determined that the rate of ATP hydrolysis by FtsA(E14R) is 60% slower than wild type FtsA under the condition tested (12 and 33 pmol min^−1^ pmol^−1^, respectively) (Fig. 2A). We suspected that introduction of a positively charged Arg at the conserved Glu near the active site could impair substrate interaction. Therefore, we measured V_max_ and K_m_ values for FtsA wild type and mutant proteins (Fig. S1C). The V_max_ values for FtsA and FtsA(E14R) were calculated to be 41.53 ± 1.76 min^−1^ and 26.51 ± 1.35 min^−1^, respectively. However, K_m_ values for FtsA and FtsA(E14R) were not significantly different (0.358 ± 0.053 mM and 0.423 ± 0.073 mM, respectively), suggesting that although FtsA(E14R) binds and hydrolyzes ATP with a similar K_m_ as wild type FtsA, it is not in a conformation associated with maximal enzymatic activity.

**Fig 2.**
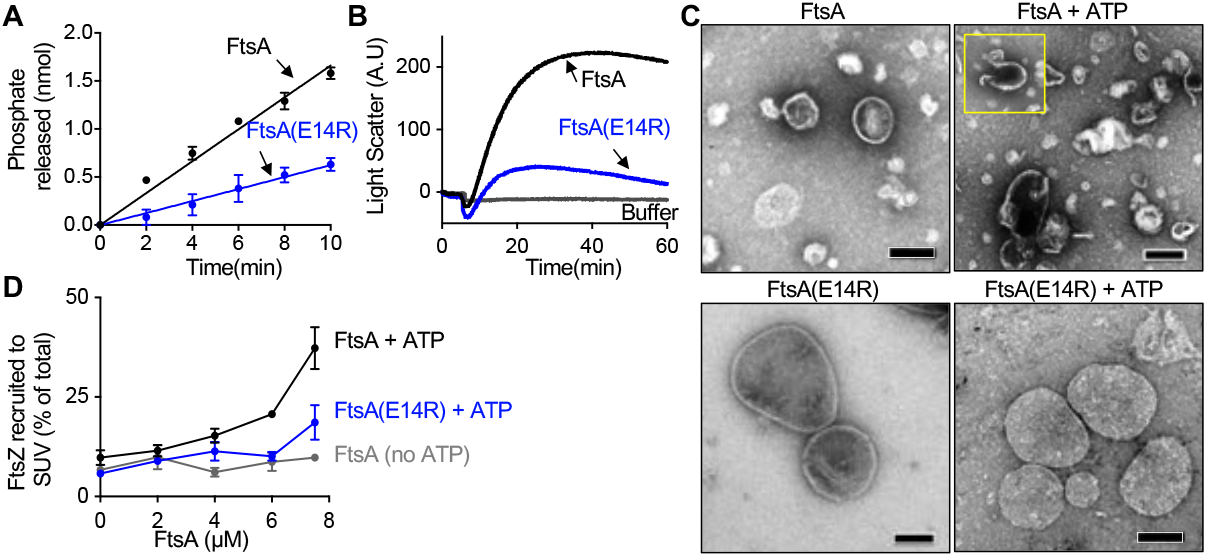
FtsA(E14R) is impaired for in vitro function. (A) Hydrolysis of ATP by FtsA (1 µM) and FtsA(E14R) (1 µM) with time as described in Materials and Methods. (B) Phospholipid reorganization by FtsA (2 µM) and FtsA(E14R) (2 µM) were monitored by 90° angle light scatter. A baseline signal was measured for 5 min and ATP was added and reactions were monitored for 60 min. (C) FtsA (4 µM) and FtsA(E14R) (4 µM) incubated in the presence and absence of ATP and visualized by TEM. Scale bars are 100 nm. Inset show additional composite image selections. (D) Phospholipid recruitment assay with reaction mixtures of FtsZ (6 µM) with GTP were pre-assembled and added to pre-incubated mixtures of FtsA or FtsA(E14R) (0-8 µM), SUV’s (250 µg ml^−1^) and ATP and fractionated by low-speed centrifugation. FtsA and FtsA(E14R) used in (A), (B), and (C) contain copurifying PLs. Data from (A) and (D) are an average of three replicates and represented as mean ± SEM and data from (B) is representative of three replicates. Pellets and supernatants for (D) were visualized by SDS-PAGE and quantified by densitometry.

Rapid ATP hydrolysis by FtsA requires phospholipids, and FtsA binds to phospholipids efficiently in vitro [7]. To confirm that FtsA(E14R) is not defective for phospholipid binding, we incubated FtsA(E14R) with SUVs (small unilamellar vesicles) prepared from *E. coli* membrane phospholipids and then collected the SUVs and bound protein by centrifugation. We determined that FtsA(E14R) was recruited to SUVs comparably to wild type FtsA in the absence and presence of ATP (Fig. S1D). We previously reported that full length FtsA copurifies with *E. coli* phospholipids, therefore we measured the amount of copurifying phospholipids with FtsA(E14R) and detected similar levels to those detected in wild type FtsA, and they are not appreciably present in FtsA(ΔMTS) preparations (Fig. S1E).

In the presence of ATP, FtsA remodels phospholipid architecture and induces tubulation of vesicles, which has been observed by TEM and 90° light scatter (LS). Tubulation of vesicles is consistent with polymerization of FtsA on the surface of a vesicle. To determine if FtsA(E14R) is defective for phospholipid remodeling (i.e., vesicle tubulation), we used LS to detect large changes in complexes after addition of ATP. Under the conditions tested, FtsA produced a large change to the LS signal in response to the addition of ATP; however, FtsA(E14R) was unable to produce a similar LS increase as wild type FtsA, which was 5-fold higher in magnitude (Fig. 2B). This suggests that FtsA(E14R) is defective for phospholipid remodeling, yet, interestingly, it binds to phospholipids and hydrolyzes ATP, although at a slower rate than wild type FtsA (Fig. 2A, S1C and S1D). Next, we directly visualized FtsA-associated vesicles by TEM to determine if protrusions and tubulations associated with lipid remodeling are produced upon incubation of FtsA(E14R) with ATP (Fig. 2C). We noted a large homogeneous population of phospholipid vesicles in reactions with FtsA(E14R); addition of ATP produced no observable effect on the architecture of the vesicles, and no rod-like tubulated structures were detected (Fig 2C). Similar to a previous report, the addition of ATP to reactions containing wild type FtsA and phospholipids resulted in large tubulated protrusions of the vesicles and short rod-like structures (Fig. 2C) [7]. These results indicate that FtsA(E14R) is unable to remodel vesicle architecture, which suggests a defect in assembly or conformational organization. Furthermore, the results are consistent with a model in which Glu 14 relays information between the adjacent active site Mg^2+^ and regions of the protein that mediate self-interaction. This is further supported by the observation that Gfp-FtsA(E14R/ΔMTS) fails to produce long self-associated structures similar to Gfp-FtsA(ΔMTS) in vivo.

Finally, to determine if FtsA(E14R) is perturbed for binding to and recruitment of FtsZ, we tested direct binding in a phospholipid recruitment assay. We performed low-speed sedimentation assays of FtsA and FtsA(E14R) with SUVs and ATP, where indicated. We observed that FtsA recruited FtsZ to phospholipids in a concentration and ATP dependent manner, demonstrating that ATP enhances FtsA recruitment of FtsZ to phospholipids, consistent with our previous report [7]. We observed that FtsA(E14R) was defective for recruitment of FtsZ to phospholipids in the presence of ATP compared to wild type FtsA (Fig. 2D). These results show that FtsA(E14R) is impaired for the interaction with FtsZ, rapid ATP hydrolysis, and ATP-dependent phospholipid remodeling.

### FtsA(ΔMTS) self-interacts with ATP to form actin-like polymers

*E. coli* FtsA(ΔMTS) is defective for ATP hydrolysis [7] (Fig. S1C), and Gfp-FtsA(ΔMTS) forms large, rod-like foci in vivo suggestive of polymer bundles [43]. Therefore, we hypothesized that FtsA(ΔMTS) may be competent for assembly of stable actin-like polymers that slowly turnover ATP and are trapped in the polymer conformation. To test this, we purified FtsA(ΔMTS) and incubated it with and without ATP, then visualized the reactions by negative staining and TEM. In the presence of FtsA(ΔMTS) and ATP, we observed long linear filaments consistent with single-stranded polymers (Fig. 3A). The polymers were highly varied in length and approximately 65 Å in width; similar to the estimated 70-80 Å width of single-stranded actin filaments visualized by TEM [50]. In the absence of ATP, we did not observe filaments, polymers, or other observable structures (Fig. 3A).

**Fig 3.**
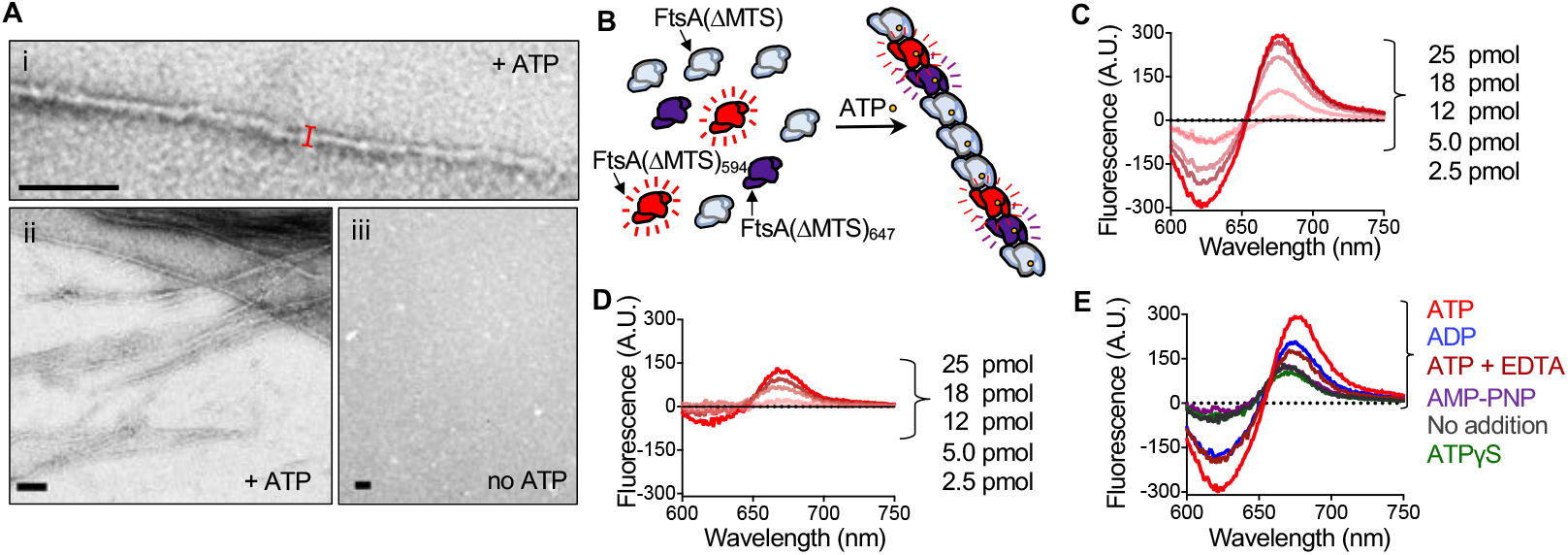
FtsA(ΔMTS) assembles into linear polymers in the presence of ATP. (A) FtsA(ΔMTS) (12 µM) incubated with (I and ii) and without (iii) ATP, and reactions were visualized by TEM as described in Materials and Methods. Scale bars are 100 nm. (B) Cartoon representation of FRET assembly reaction. AF594 labeled FtsA(ΔMTS) (donor), AF647 labeled FtsA(ΔMTS) (acceptor) was mixed with FtsA(ΔMTS) and reactions were excited at the donor excitation wavelength and scanned for the emission of each fluorophore. Close proximity of the donor fluorophore to the acceptor fluorophore causes a large increase in acceptor emission fluorescence, as occurs by the addition of ATP. (C) FRET using increasing concentrations of Alexa Fluor 647 labeled FtsA(ΔMTS) (2.5 to 25 pmol) were mixed with a fixed concentration of Alexa Fluor 594 labeled FtsA(ΔMTS) (100 pmol) and FtsA(ΔMTS) for a total of 600 pmol. Reactions were monitored after addition of ATP as described in Materials and Methods. (D) FRET reactions were assembled and monitored as in (C) but without the addition of ATP. (E) FRET emission if Alexa Fluor 647 labeled FtsA(ΔMTS) (25 pmol) mixed with Alexa Fluor 594 labeled FtsA(ΔMTS) (100 pmol) and FtsA(ΔMTS) (475 pmol) was monitored after addition of ATP, ADP, ATP with EDTA, ATPγS, or no nucleotide addition.

Next, we developed a fluorescence-based self-assembly assay to more quantitatively compare FtsA(ΔMTS)-FtsA(ΔMTS) interactions. We labeled two populations of FtsA(ΔMTS), one with Alexa fluor 594 (AF_594_; donor) and another with Alexa fluor 647 (AF_647_; acceptor), to measure energy transfer from donor to acceptor fluorophore during ATP-dependent assembly (Figs. 3B and S2A). We incubated donor-labeled FtsA(ΔMTS) with acceptor-labeled FtsA(ΔMTS) and observed a large emission signal in the presence of ATP (Fig. 3C), along with a corresponding decrease in donor-emission near 625 nm. The emission amplitudes that we measured titrated with the amount of acceptor-labeled FtsA(ΔMTS) in each reaction. We repeated this experiment in the absence of ATP and detected low level emission of the acceptor-labeled FtsA(ΔMTS), even at high acceptor concentration, compared to the emission observed in the presence of ATP (Figs. 3D and S2B). Together, these results demonstrate that ATP enables the energy transfer between donor- and acceptor-labeled populations of FtsA(ΔMTS), consistent with FtsA(ΔMTS) self-interaction occurring under conditions that promote polymerization as detected by TEM (Fig. 3A).

Next, we compared the acceptor FRET signal associated with various nucleotide analogs and observed that ATP supported energy transfer most efficiently (Fig. 3E). The ATP-dependent emission signal was reduced in the presence of EDTA, which would prevent Mg^2+^ from binding in the active site, and the emission was also lower with ADP, compared to ATP. Neither ATPγS nor AMP-PNP supported energy transfer over the nucleotide-free control (Fig. 3E). Next, if ATP induces the assembly of stable FtsA(ΔMTS) polymers, then we hypothesized that we would be able to collect polymers by high-speed centrifugation. We incubated increasing amounts of FtsA(ΔMTS) with and without ATP, collected pellet and supernatant fractions after centrifugation at 160,000 x *g* for 30 min, and analyzed both fractions by SDS-PAGE. We observed that FtsA(ΔMTS) fractionated predominantly with the pellet (>50%) in the presence of ATP, and predominantly with the supernatant (>95 %) when ATP was omitted (Fig. S2C and Fig. S2D). Moreover, the relative amount of FtsA(ΔMTS) in the pellet increased with the total FtsA(ΔMTS) concentration in the reaction, suggesting that assembly efficiency increases with FtsA(ΔMTS) concentration. We compared the relative abilities of several nucleotides to support polymerization of FtsA(ΔMTS) in the sedimentation assay and observed that ATP yielded the largest amount of pellet-associated FtsA(ΔMTS) (50.0%), and this amount was reduced by EDTA (16.5%) (Fig. S2E) or with ADP (22.4%), ATPγS (19.5%), or AMPPNP (<5%) (Fig. S2E). Finally, we utilized 90° light scatter to compare polymerization efficiency of FtsA(ΔMTS) at different temperatures, protein concentrations and nucleotides. We observed that at 37 °C, addition of ATP produced a rapid increase in LS, which plateaued after approximately 10 min, and the amplitude change upon addition of ATP was reduced at lower temperatures (30 °C and 23 °C) (Fig. S2F). The amplitude of the light scatter change with ATP also increased with FtsA(ΔMTS) concentration up to 20 μM (Fig. S2G). Finally, the addition of ADP to FtsA(ΔMTS) also led to a modest increase in LS amplitude; however, it was ∼60% lower than the increase obtained with ATP (Fig. S2H), which is consistent with results from sedimentation assays (Fig. S2E). Addition of EDTA to the assembly reactions prevented the amplitude increase associated with both ATP and ADP, indicating that Mg^2+^ association is likely important for polymerization (Fig. S2H). Together, these results demonstrate that FtsA(ΔMTS) self-interacts in the presence of ATP and Mg^2+^ to form actin-like polymers.

### FtsA(ΔMTS) polymers bind directly to FtsZ polymers but are destabilized by non-polymerized FtsZ

To determine if polymers of FtsA(ΔMTS) stably interact with polymers of FtsZ, we developed a bead recruitment assay to overcome limitations associated with interpreting cosedimentation results of two polymer subassemblies. In this assay, we pre-assembled GMPCPP-stabilized polymers of FtsZ, fused to Gfp and containing a six-histidine tag (H_6_-Gfp-FtsZ), then recruited the polymers to the surface of a cobalt bead, which has an average diameter of 55 μm (Fig. 4A). H_6_-Gfp-FtsZ has previously been reported to hydrolyze GTP and polymerize in vitro, similar to wild type FtsZ [51]. Next, we pre-assembled FtsA(ΔMTS), labeled with AF 594, with ATP to induce polymerization and then tested for recruitment to cobalt beads in the presence and absence of FtsZ and GMPCPP (Fig. 4A). We observed bundles of FtsZ and FtsA(ΔMTS) fluorescence in the presence of nucleotides GMPCPP and ATP, respectively, but no localized fluorescence in the absence of nucleotides (Fig. 4Ai, S3A and S3B). Moreover, H_6_-Gfp-FtsZ localized to the bead surface (Fig. 4Aii). Fluorescence microscopy does not resolve single-stranded polymers, but does distinguish soluble protein, as diffuse fluorescence, from foci, which is suggestive of higher order assembly architecture. Incubation of pre-assembled H_6_-Gfp-FtsZ polymers with FtsA(ΔMTS) polymers resulted in colocalization of FtsA(ΔMTS) to the bead surface in the presence of His_6_-Gfp-FtsZ (i), but not when H_6_-Gfp-FtsZ was omitted (Fig. 4Ai and 4Aiii). Therefore, under conditions that promote polymerization of both proteins (i.e., with nucleotides GMPCPP and ATP), H_6_-Gfp-FtsZ recruits FtsA(ΔMTS) to the bead surface, suggesting that both polymer subassemblies interact directly.

**Fig 4.**
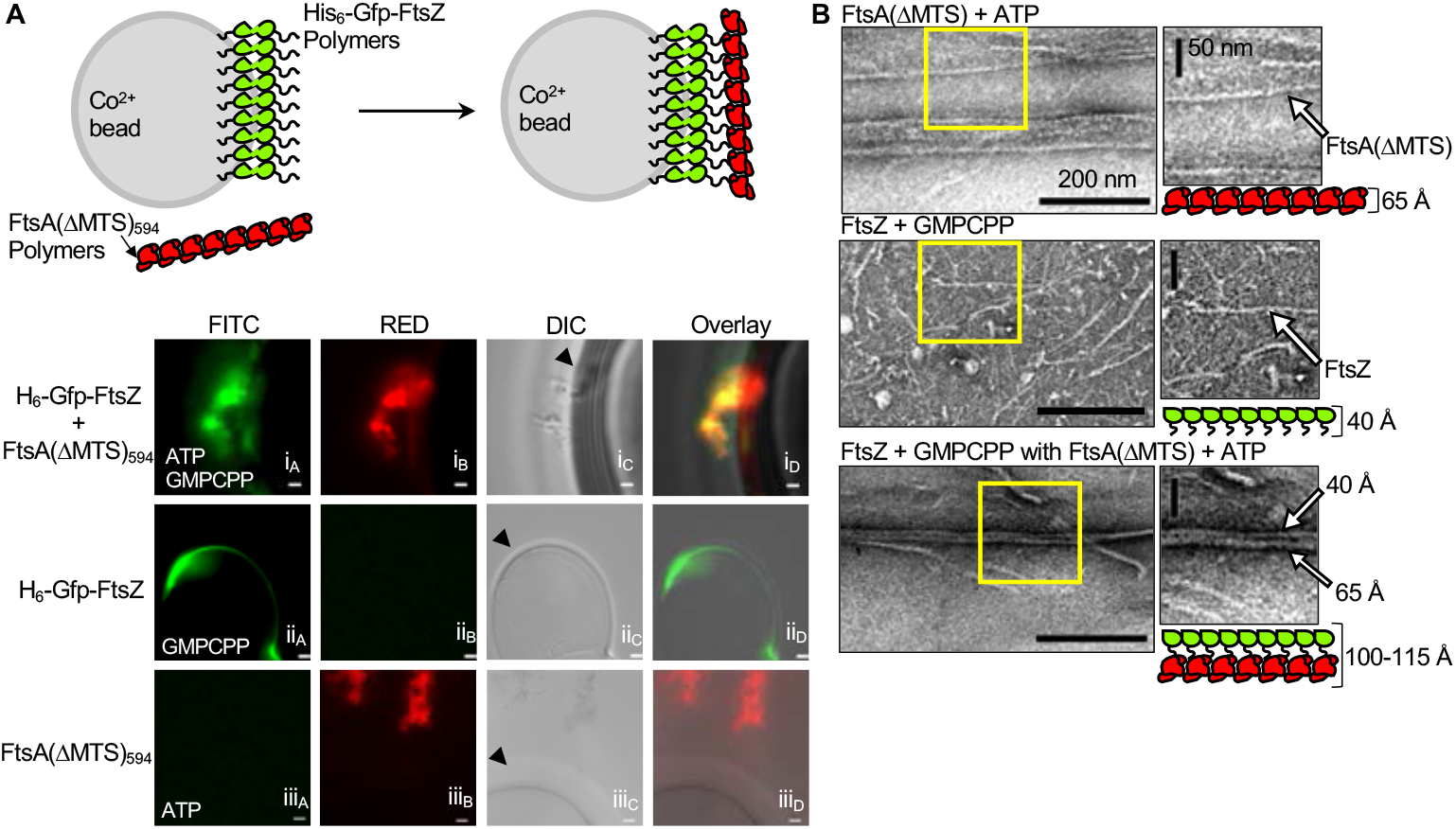
Coassembly of FtsA(ΔMTS) and FtsZ polymers. (A) Upper panel shows cartoon representation of recruitment of FtsA(ΔMTS)_594_ polymers by His_6_-Gfp-FtsZ complexes on cobalt beads. Fluorescence and DIC microscopy of reaction mixtures of (i) H_6_-Gfp-FtsZ(green), FtsZ, GMPCPP, and cobalt beads, added to reactions of AF594 FtsA(ΔMTS)(red), FtsA(ΔMTS) and ATP; (ii) H_6_-Gfp-FtsZ(green), FtsZ, GMPCPP, and cobalt bead (iii) AF594 FtsA(ΔMTS)(red), FtsA(ΔMTS), ATP, and beads were assembled and visualized as described in Materials and Methods. Scale bars are 1 µm for (i) and 5 µm for (ii) and (iii). (B) FtsZ (8 µM) and GMPCPP, FtsA(ΔMTS) (8 µM) and ATP, and FtsZ (8 µM) and GMPCPP added to pre-assembled reaction mixtures of FtsA(ΔMTS) (8 µM) and ATP were visualized by TEM as described in Materials and Methods. Yellow box highlights areas enlarged for detail.

To elucidate the in vitro protein architecture in mixtures of FtsA(ΔMTS) polymers, assembled with ATP, and FtsZ polymers, assembled with GMPCPP, we visualized polymers alone and together by TEM. We observed that GMPCPP-stabilized FtsZ polymers were long, single-stranded and approximately 40-45 Å in width, consistent with previous reports [7] (Fig. 4B). In contrast, FtsA(ΔMTS) polymers were visibly wider, approximately 60-75 Å in width, and also long and single-stranded (Fig. 3A and 4B). When FtsA(ΔMTS) and FtsZ polymers were incubated together and then visualized by TEM, we observed wide filaments (100-120 Å) and a population of polymers with a distinct paired-filament morphology. These doublet polymers also had a width of approximately 100-120 Å (Fig. 4B) and were composed of two aligned polymers of widths (∼60 Å and 40 Å) (Fig. 4B). The morphological features of the paired filament are consistent with an FtsZ polymer aligned with and bound to an FtsA(ΔMTS) polymer.

Next, we interrogated if FtsZ has a direct effect on the propensity for or degree of FtsA polymerization. Direct visualization of polymers is non-quantitative, therefore, to test if FtsZ, with or without GMPCPP or GTP, modifies ATP-dependent assembly of FtsA(ΔMTS), we used the fluorescence-based FtsA(ΔMTS) subunit interaction FRET assay. First, we tested if FtsZ, assembled with GMPCPP, modified the acceptor emission, which would suggest that FtsZ modifies the self-interaction of FtsA(ΔMTS). We observed no decrease in fluorescence emission by acceptor-labeled FtsA(ΔMTS) in the presence of ATP, FtsZ (8 μM) and GMPCPP (Fig. 5A). Similarly, we also observed no effect when FtsZ and GTP were added to the reaction instead of GMPCPP (Fig. 5A). However, when GMPCPP or GTP was omitted, we observed that addition of FtsZ (8 μM) reduced the fluorescence emission of acceptor-labeled FtsA(ΔMTS) by ∼40% under the conditions tested and in a concentration dependent manner (Fig. 5A and S3C).

**Fig 5.**
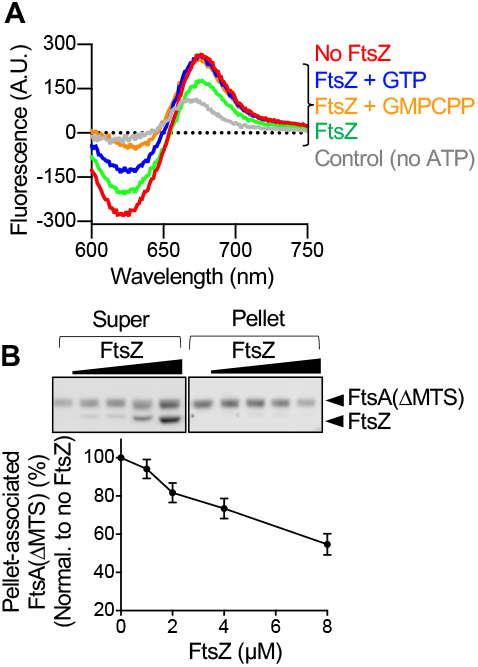
FtsA(ΔMTS) polymers are regulated by FtsZ. (A) FRET emission of reactions without FtsZ (red) of with FtsZ (8 µM) mixed with Alexa Fluor 647 labeled FtsA(ΔMTS) (25 pmol), Alexa Fluor 594 labeled FtsA(ΔMTS) (100 pmol) and FtsA(ΔMTS) (475 pmol) and monitored after addition of ATP as described in Materials and Methods. Where indicated, FtsZ was pre-assembled with GTP (blue) or GMPCPP (gold), or incubated without nucleotide (green). FtsA(ΔMTS) without ATP is shown as a control (gray). Data is representative of three independent replicates. (B) FtsZ (0 to 8 µM) was added to pre-assembled reaction mixtures of FtsA(ΔMTS) (8 µM) and ATP, incubated for 5 min, fractionated by ultracentrifugation, and analyzed by SDS-PAGE and densitometry. and Graph shown displays an average of three replicates represented as mean ± SEM.

The reduced FtsA(ΔMTS) acceptor fluorescence in the presence of ATP and non-polymerized FtsZ suggests that FtsZ destabilizes the FtsA(ΔMTS) self-interaction, which would lead to disassembly of FtsA(ΔMTS) polymers. To test this directly, we performed high speed sedimentation assays to collect ATP-stimulated FtsA(ΔMTS) polymers and titrated FtsZ concentration. First, we pre-assembled FtsA(ΔMTS) (8 μM) with ATP (4 mM) and then added increasing concentrations of FtsZ (0 to 8 μM). After incubation for 5 min, FtsA(ΔMTS) polymers were collected by high-speed centrifugation, and then supernatants and pellets were analyzed by SDS-PAGE for the presence of FtsA(ΔMTS) and FtsZ. We observed that although we titrated the amount of FtsZ in the reaction, we failed to recover FtsZ in the sedimentation pellet with FtsA(ΔMTS) and ATP (Fig. 5B). In addition, the amount of FtsA(ΔMTS) fractionating with the pellets decreased as the FtsZ concentration in the reaction increased (Fig. 5B), suggesting that FtsZ, without GTP, destabilizes FtsA(ΔMTS) polymers assembled with ATP in agreement with the FRET result (Fig. 5A). Together, these results show that unassembled FtsZ destabilizes FtsA(ΔMTS) polymers, and suggest that stable FtsZ polymers and FtsA(ΔMTS) polymers align together. This suggests a biochemical model in which FtsZ polymers facilitate FtsA polymerization, and depolymerized FtsZ facilitates FtsA depolymerization.

### MinC disrupts FtsA-FtsZ complexes releasing FtsZ from the membrane

According to a Z-ring model in which FtsZ and FtsA function together as coregulated, co-assembled polymers, FtsZ polymerization inhibitors, such as MinC, should destabilize FtsZ polymers, thereby disrupting the interaction between FtsA and FtsZ that occurs at the membrane. Disassembly activity would be further exacerbated if the recognition regions on FtsZ overlapped. MinC promotes disassembly of FtsZ polymers in vitro and both proteins, MinC and FtsA, bind near the C-terminal end of FtsZ [7, 52]. To test if MinC disrupts the interaction of polymerized FtsZ with FtsA-associated SUVs, we performed phospholipid recruitment assays. First we incubated FtsZ (3 μM) with GTP (2 mM), to induce polymerization, and where indicated, MinC (5 μM), to destabilize FtsZ polymers. Then, we added FtsA-associated liposomes containing 3 μM FtsA and 250 μg ml^−1^ SUVs to the reactions containing FtsZ, in the absence and presence of MinC, and collected SUV-protein complexes by centrifugation. Proteins in supernatant and PL pellet fractions were visualized by SDS-PAGE and Coomassie staining. We observed that when either GTP or FtsA were omitted from the reaction, FtsZ remained in the supernatant, not associated with SUVs (Fig. 6A). However, in reactions containing FtsZ, GTP and FtsA, 85.6% of FtsZ fractionated with FtsA-associated SUVs in the pellet. When FtsZ was incubated with GTP and MinC, then recruited to SUVs with FtsA, only 24.6% of FtsZ fractionated with the pellet. In addition to destabilizing FtsZ polymers, these results demonstrate that MinC prevents recruitment of FtsZ polymers to PL-associated FtsA.

**Fig 6.**
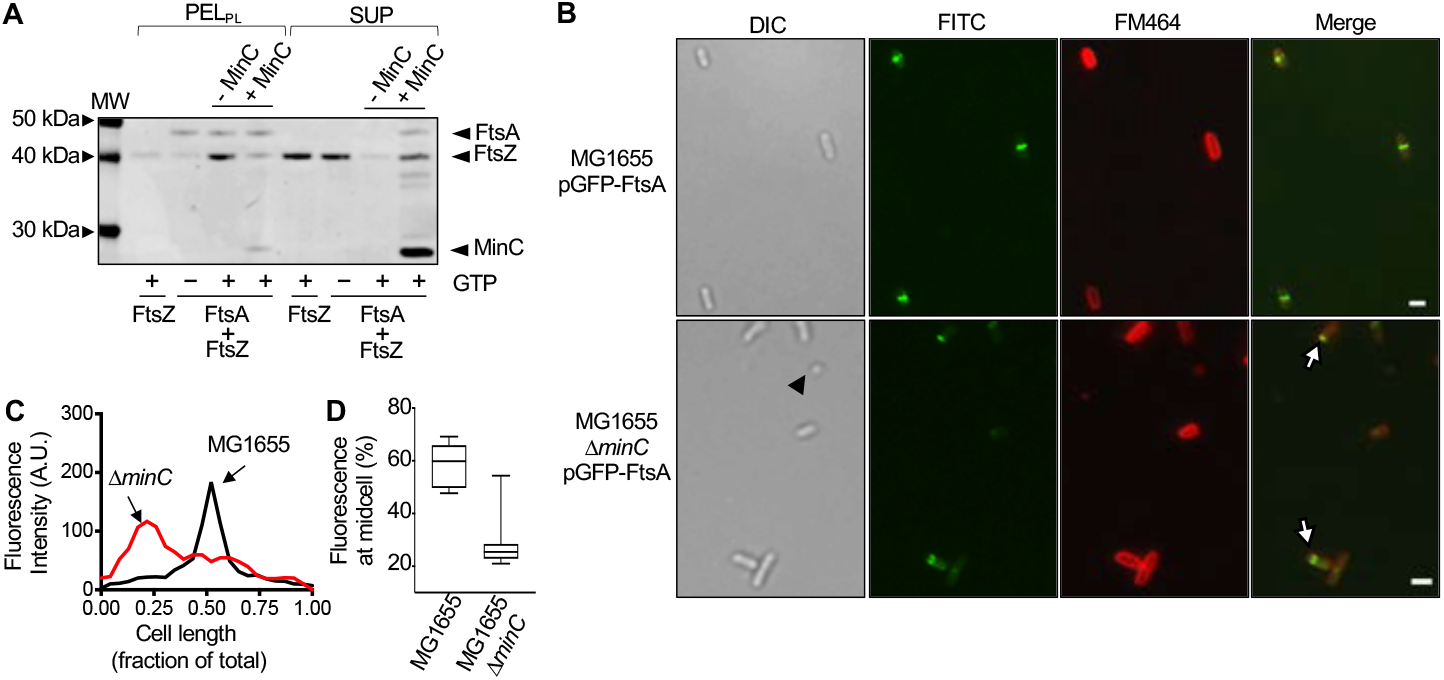
Mislocalization of Gfp-FtsA in *minC* deletion strain suggests regulation by FtsZ. (A) Phospholipid recruitment assay to monitor recruitment of FtsZ to FtsA-associated SUVs. Reaction mixtures containing FtsZ (3 µM) with and without MinC (5 µM) were incubated with GTP, where indicated, for 2 min. FtsA (2 µM), SUVs (250 µg ml^−^ _1_), and ATP were added, and then reactions were incubated for an additional 10 min. Reactions were fractionated by low-speed centrifugation and PL-associated pellets and supernatants were visualized by SDS-PAGE. (B) Fluorescence microscopy and differential interference contrast (DIC) microscopy of *E. coli* MG1655 and MG1655 Δ*minC* cells expressing Gfp-FtsA (green) and stained with FM-464 (red). Scale bars are 2 µm. Black arrow indicates a minicell. White arrows indicate polar Gfp-FtsA rings. (C) Fluorescence intensity across the long axis of the cell was measured and plotted to determine the relative localization of Gfp-FtsA for *E. coli* MG1655 and MG1655 *ΔminC* cells. (D) Box and whiskers plot showing percent fluorescence at midcell of *E. coli* MG1655 and MG1655 *ΔminC* cells expressing Gfp-FtsA (n=10). The extent of the box encompasses the interquartile range of the fluorescence intensity, whiskers extend to the maximum and minimum fluorescence intensities, and the line within each box represents the median.

In a dividing cell, if FtsZ and FtsA function together as co-regulated, co-assembled polymers, then the location and assembly state of FtsZ could therefore predict, and actively regulate, where FtsA assembles. In wild type cells, Gfp-FtsA localizes to Z-rings with high efficiency (Fig. 1B). We would further predict that modification of the normal distribution of FtsZ polymers in vivo would also lead to a consequent change in FtsA distribution. To test this in vivo, we deleted the gene encoding MinC, which exhibits a time-averaged localization near the cell poles, and monitored Gfp-FtsA localization. Deletion of *minC* leads to polar ring formation and minicells when the polar rings complete division [53]. Thus, the presence of an excess of polymerized FtsZ at multiple locations, leads to division rings that complete septation at multiple locations. This could only proceed if the rest of the cell division proteins also assembled at those sites. Consistent with this, overexpression of FtsZ also leads to excess Z-rings and polar division events, in addition to filamentous cells. Consequently, disruption of MinC activity, leading to aberrant FtsZ polymerization and localization must also lead to aberrant FtsA localization if FtsA is required to complete the polar septation event. To determine if deletion of *minC* leads to aberrant FtsA distribution in the cell, we expressed Gfp-FtsA from a plasmid in wild type (MG1655) and *minC* deletion strains and then monitored fluorescence as a function of cell length (Fig. 6B and 6C). We observed that in wild type cells expressing Gfp-FtsA, approximately 60% of the total cellular fluorescence was located at the midcell region and Z-rings were present (Figs. 6B, 6C and 6D). In contrast, in cells deleted for *minC*, approximately 25% of the total cellular fluorescence was present at midcell, and polar Z-rings were detected in many cells. In a *minC* deletion background, ClpX contributes to the regulation and function of active Z-rings, since the additional deletion of *clpX* causes severe filamentation and exacerbates aberrant Z-ring placement and minicell formation [54]. We monitored Gfp-FtsA localization in the *minC clpX* double deletion strain and observed polar Gfp-FtsA rings, similar to cells deleted for *minC* (Fig. S4A). We also tested if deletion of *slmA* also leads to mislocalized Gfp-FtsA. In addition to MinC, SlmA is a known FtsZ polymer destabilizing protein that binds to the nucleoid and prevents Z-ring formation over the nucleoid [55]. Expression of Gfp-FtsA in a *E. coli slmA* deletion strain showed ring assembly at midcell and in locations where the nucleoids were beginning to appear segregated (Fig. S4B). Consistent with previous reports, cells deleted for *slmA* do not show major changes in longitudinal Z-ring placement [55]. Together, our results show that the FtsZ polymerization inhibitor MinC, prevents FtsA from recruiting FtsZ polymers to phospholipids and, in the absence of MinC, mislocalized FtsZ polymers that are present in the cell are sufficient to cause mislocalization of FtsA.

## Discussion

Here, we demonstrate that a truncated *E. coli* FtsA variant, FtsA(ΔMTS), forms linear ATP-dependent polymers in vitro that are clearly observable by TEM (Fig. 3A) and readily detectable in direct biophysical assays (Fig. 3C, S2B-H). Most surprisingly, we observed that FtsZ induces disassembly of FtsA(ΔMTS) polymers, but only in the absence of GTP (Fig. 5A, 5B and S3C). In contrast, when FtsZ polymerizes, either with GTP or GMPCPP, this prevents the disassembly and, as we show with the latter nucleotide, FtsZ polymers directly associate with FtsA(ΔMTS) polymers. These results suggest a model by which FtsA and FtsZ copolymers laterally associate at the membrane. This is consistent with the suggestion that in vivo, FtsA and FtsZ polymers assemble at midcell together and exhibit treadmilling [56]. It has been further suggested that FtsZ-interacting proteins, such as ZapA, modify the spatial order and dynamics of membrane associated FtsZ bundles, but do not directly affect treadmilling [57]. Here, we observed that MinC displaces FtsZ and prevents FtsA from recruiting FtsZ to SUVs, suggesting that this activity underlies the potent disruption of polar Z-rings when MinC is present in the cell and the resulting minicell phenotype that occurs when *minC* is deleted from cells. During normal division, only 30% of FtsZ localizes to the division site, leaving 70% of FtsZ remaining in the cytoplasm, but that 70% does not produce division events, likely because the assembly state is maintained in a largely non-polymerized capacity by regulators of FtsZ assembly positioned throughout the cell.

While phospholipid engagement is essential for rapid ATP hydrolysis and nucleotide cycling by FtsA in vitro, the precise steps of nucleotide binding, hydrolysis and release, and how those steps regulate FtsA conformation, remain to be determined. Interestingly, substitution of Glu 14 with Arg, which is adjacent to the Mg^2+^ in the model of the active site, does not completely prevent ATPase activity (Fig. 2A and S1C), suggesting that, while Glu 14 plays a role, it is not the major catalytic residue involved in hydrolysis of the ATP gamma phosphate. FtsA(E14R) is defective for recruiting FtsZ polymers to SUVs in vitro and fails to form rings in vivo. Although FtsA(E14R) binds to SUV’s, in the presence of ATP it does not induce tubulation, a hallmark of polymerization, and introduction of the E14R mutation in Gfp-FtsA(ΔMTS) prevents the accumulation of cytoplasmic foci in cells. These results have several implications. First, ATP hydrolysis does not appear to absolutely require polymerization, which is reminiscent of differing hydrolysis rates of G-actin (globular) and F-actin (filamentous) [58] and in contrast to FtsZ, where residues from adjacent protomers are required for enzymatic activity at a dimer interface [59]. If FtsA(E14R) can engage PLs and hydrolyze ATP, then why is it defective for polymerization? It is likely that Glu 14 regulates and/or stabilizes a key conformational transition that facilitates polymerization and direct FtsZ interaction, which must also be coordinated with PL engagement. This implies that FtsA therefore undergoes multiple conformational transitions, with full activity elicited by engagement of both PLs and ATP.

In our model for FtsA-FtsZ interactions (Fig. 7A and 7B), FtsA binds directly to PLs in the absence and presence of ATP by insertion of the C-terminal MTS into the bilayer (Fig. S1D) [7]. When PL-associated FtsA binds ATP, this induces polymerization leading to localized membrane distortion and tubulation. When FtsZ is present, FtsZ binds to FtsA-associated phospholipids in a reaction that is stimulated with GTP. Polymerized FtsZ is capable of forming copolymers with FtsA, and this is likely the conformation that leads to productive division. Without GTP, FtsZ destabilizes FtsA polymers, leading to active disassembly of transient polymeric subcomplexes. This effect is reciprocal, as it was previously reported that FtsA can also induce disassembly of FtsZ polymers [7]. This reciprocal effect establishes that FtsA and FtsZ polymers are coordinately regulated and their assembly state likely modifies the commitment to the later steps of division. We predict that the next step after engagement of FtsZ polymers by FtsA is activation of FtsN; however, it is not currently known how this would occur. One possibility is that the localized concentration of FtsA may recruit a critical mass of FtsN to a single location to initiate septal PG synthesis on the periplasmic side of the where FtsA-FtsZ complexes assemble (Fig. 7B). Both FtsA and FtsN appear to be critical for recruiting and activating cell wall synthetases [60]. There are two major advantages to this model: (i) establishment of the Z-ring would be responsive to the actions of FtsZ assembly regulators inside the cell, such as MinC, SlmA, and OpgH [61], and (ii) progression through the cell cycle would also be responsive to ATP and GTP levels. Notably, Z-rings have been shown to rapidly disassemble when ATP is depleted, indicating that cellular energy availability also regulates assembly of division complexes [42].

**Fig 7.**
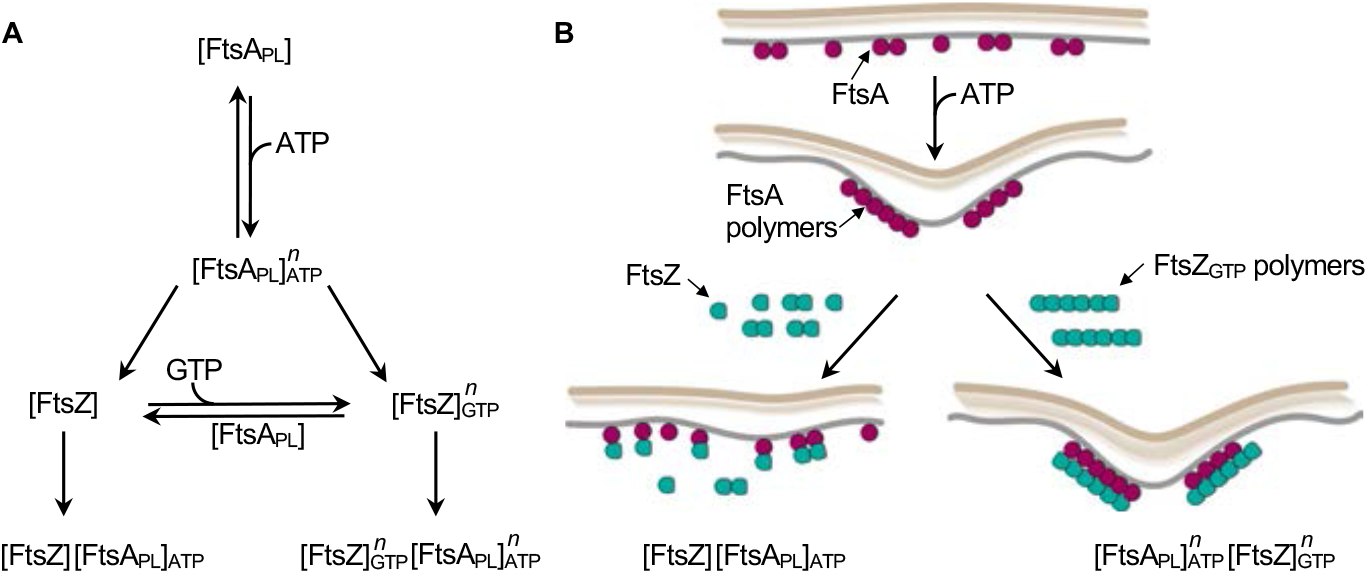
Model of FtsA-FtsZ polymer coregulation during *E. coli* cell division. (A) Phospholipid bound FtsA [FtsA _PL_] binds to ATP, which promotes polymerization [FtsA PL]^n^_ATP_ on the membrane. In the presence of FtsZ polymers [FtsZ]^n^, FtsA polymers coassemble with FtsZ polymers ([FtsZ]^n^_GTP_ [FtsA]^n^_ATP_) on the membrane surface, thus localizing the FtsZ polymers to the membrane at midcell during Z-ring formation. Conversely FtsZ monomers [FtsZ], which may be produced by the action of FtsZ-assembly regulators, such as MinC, or by low GTP levels, antagonize FtsA polymers inducing disassembly. This prevents establishment of nascent Z-rings containing FtsZ and FtsA and premature activation of cell wall synthetases. (B) Cartoon representation of the model for cell division in (A). Briefly, FtsA polymers assemble with ATP and recruit FtsZ polymers to the membrane. The local deformation of membrane architecture leads to FtsA-mediated recruitment and activation of FtsN and downstream cell division protein for PG synthesis.

In eukaryotes, there are a variety of actin-modifying proteins that regulate the assembly state and functional roles of actin in cells; these include actin severing and capping proteins, bundling and branching proteins, as well as a nucleators, such as the formin Bni1p, which is involved in polarized cell growth in *Saccharomyces cerevisiae* [62]. Several proteins have been suggested to interact directly with FtsA to potentially modify the polymerization state, including ZipA, FtsN, and FtsEX [14, 16, 17, 45, 63]; all have been suggested to destabilize or bypass FtsA-FtsA interactions. Interestingly, FtsA is highly conserved indicating that the ATP hydrolysis and polymerization functions of FtsA have been evolutionarily preserved and are therefore critical for function [46]. However, it is possible that the assembly state of FtsA must be fine-tuned to both respond to the assembly state of FtsZ and recruit the appropriate level of cell wall synthetases to the septum. A previous model suggested that FtsA is in competition for interactions with itself and downstream division proteins, and this equilibrium may be modified by interacting proteins, such as ZipA [43, 44]. Polymerization of FtsA may be required to correctly recruit FtsZ but must be limited by other factors to prevent premature assembly and activation of the PG synthetases. Given the abundance of actin-regulating proteins in eukaryotes, it will be interesting to determine how many bacterial cell division proteins engage and modify the assembly state of FtsA, ATP hydrolysis, coordinated regulation with FtsZ polymers, and activation of cell wall synthesis proteins.

## Materials and Methods

### Bacterial strains and plasmid construction

The strains and plasmids used in this study are listed in Table 1. The *ftsA* gene was amplified from pET-*ftsA* and cloned into pQE9 using *BamHI* and an engineered *NdeI* restriction site. Site-specific mutants in *ftsA* were constructed by site-directed mutagenesis of *ftsA* expression plasmids using the QuikChange II XL mutagenesis system (Agilent) and confirmed by sequencing. pQE-9-*ftsA* was mutagenized to create FtsA(E14R), FtsA(ΔMTS), and FtsA(E14R/ΔMTS). FtsA(ΔMTS) was created by inserting a TAA stop codon at amino acid position 405 in *ftsA*. Plasmid pSEB293 [4], containing *gfp-ftsA*, was mutagenized to create Gfp-FtsA(E14R), Gfp-FtsA(ΔMTS), Gfp-FtsA(E14R), and Gfp-FtsA(E14R/ΔMTS). pET-*ftsA* was mutagenized to create FtsA(E14R) and FtsA(ΔMTS).

**Table 1.** *E. coli* strains and plasmids

| Strain or plasmid | Relevant Genotype | Source, reference, or construction |
| --- | --- | --- |
| **Strain** |  |  |
| MG1655 | LAM- *-rph1* | [67] |
| BL21(λde3) | *F- ompT gal dcm lon hsdSB(rB- mB-)*<br>*$\lambda$(de3[*lacI lacUV5-T7 gene 1 lnd1 sam7 nin5*])* | EMD Millipore, USA |
| JC0232 | MG1655 (Δ*minC::frt*) | [54] |
| JC0259 | MG1655 (Δ*clpX::kan*) | [54] |
| JC0292 | MG1655 (Δ*minC::frt* Δ*clpX::frt*) | [54] |
| JC0390 | MG1655 (Δ*araEp:kan* PCP<sub>18</sub>-*araE*) | [51] |
| MV0196 | MG1655 (Δ*slmA::frt*) | [51] |
| MCA12 | *F-*, [*araD139*]<sub>B/r</sub>, *leu-260::Tn10*, *ftsA12(ts)*,<br>*$\Delta$(argF-lac)169*, $\lambda^-$, *e14-*, *flhD5301*, *$\Delta$(fruK-yeiR)*<br>*725(fruA25)*, *relA1*, *rpsL150(strR)*, *rbsR22*,<br>*$\Delta$(fimB-fimE)632(::IS1)*, *deoC1* | [49] |
| MCA27 | *F-*, [*araD139*]<sub>B/r</sub>, *leu-260::Tn10*, *ftsA27(ts)*,<br>*$\Delta$(argF-lac)169*, $\lambda^-$, *e14-*, *flhD5301*, *$\Delta$(fruK-yeiR)*<br>*725(fruA25)*, *relA1*, *rpsL150(strR)*, *rbsR22*,<br>*$\Delta$(fimB-fimE)632(::IS1)*, *deoC1* | [49] |
| **Plasmid** |  |  |
| pET-24b | *kan*, P<sub>T7</sub> promoter | EMD Millipore, USA |
| pET-*ftsA* | *kan* | [7] |
| pET-*ftsA*(ΔMTS) | *kan* | [7] |
| pET-*ftsA*(E14R) | *kan* | This study |
| pET-*ftsZ* | *kan* | [65] |
| pET-*minC* | *kan* | [66] |
| pET-H<sub>6</sub>-Gfp-*ftsZ* | *kan* | [51] |
| pSEB293 | *amp* P<sub>ara</sub>::Gfp-*ftsA* | [4, 33] |
| pBAD-Gfp-*ftsA*(E14R) | *amp* P<sub>ara</sub>::Gfp-*ftsA*(E14R) | This study |
| pBAD-Gfp-*ftsA*(ΔMTS) | *amp* P<sub>ara</sub>::Gfp-*ftsA*(ΔMTS) | This study |
| pBAD-Gfp-*ftsA*(ΔMTS)(E14R) | *amp* P<sub>ara</sub>::Gfp-*ftsA*(ΔMTS)(E14R) | This study |
| pQE9 | *amp* P<sub>T5</sub> promoter | [68] |
| pQE9-*ftsA* | *amp* | This study |
| pQE9-*ftsA*(E14R) | *amp* | This study |
| pQE9-*ftsA*(ΔMTS) | *amp* | This study |
| pQE9-*ftsA*(ΔMTS)(E14R) | *amp* | This study |

### Protein purification and ATP hydrolysis assays

FtsA, FtsA(E14R), and FtsA(ΔMTS) were purified as previously reported [7]. FtsZ and H_6_-Gfp-FtsZ were purified as previously reported [64, 65]. MinC was purified as previously described [66]. ATP hydrolysis assays were performed by measuring the amount of inorganic phosphate released at 37°C in reaction mixtures (25 µl) with a reaction buffer containing 50 mM Tris-HCl (pH 7.5), 150 mM KCl, and 10 mM MgCl_2_, FtsA, FtsA(E14R), or FtsA(ΔMTS) (1 µM) and ATP (4 mM) using Biomol Green (Enzo Life Sciences) and comparing to a phosphate standard curve. Protein concentrations refer to FtsA monomers, FtsZ monomers and MinC dimers.

### Transmission electron microscopy

Reactions of reaction buffer, FtsA(ΔMTS) (12 µM), FtsA (4 µM), FtsA(E14R) (4 µM), and ATP (4 mM), where indicated, were incubated for 5 min at 23°C, applied to a 300-mesh carbon/formvar coated grid, fixed with glutaraldehyde (2.5%) and negatively stained with uranyl acetate (2%). Samples were imaged by transmission electron microscopy using a JEM-2100 80 Kev instrument.

### FRET assays

Several Cys residues in FtsA(ΔMTS) are surface exposed when mapped on the structural model of *E. coli* FtsA (Fig. S2A). Two populations of FtsA(ΔMTS) were labeled with Alexa Fluor 594 (donor) and Alexa Fluor 647 (acceptor) according to the manufacturer’s protocol (ThermoFisher) to a degree of labeling of 0.5-2.0. This widely used donor-acceptor pair has a Förster radius of 79 Å, which corresponds closely to a distance equivalent to two adjacent FtsA protomers, with each protomer being approximately 40 Å in diameter. To monitor energy transfer, reaction mixtures (80 µl) were incubated in an assembly buffer containing 50 mM Tris-HCl (pH 7.5), 150 mM KCl, 4 mM MgCl_2_, and, where indicated, AF594-labeled FtsA(ΔMTS) (100 pmol), AF647-labeled FtsA(ΔMTS) (0 to 25 pmol), FtsA(ΔMTS) (475 to 500 pmol), FtsZ (4 to 8 µM), and EDTA (15 mM) for 5 min at 37°C. Fluorescence emission was scanned from 600 to 800 nm at an excitation wavelength of 590 nm. To compare nucleotides, ATP (4 mM), ADP (4 mM), or ATPγS (4 mM), were added, where indicated, and reactions were incubated at 37°C for 10 min, then emission spectra were collected. To correct for background and off-peak non-specific donor signal, equivalent reactions omitting acceptor fluorophore were scanned and curves were subtracted.

### Assembly and recruitment assays

To measure FtsA(ΔMTS) polymer formation by ultracentrifugation, reaction mixtures (25 µl) containing assembly buffer, FtsA(ΔMTS) (0 to 16 µM), and ATP (4 mM), where indicated, were assembled. Reactions were incubated at 23 °C for 10 min and, where indicated, FtsZ (0 to 8 µM), or FtsZ (0 to 8 µM) preassembled with GTP (4 mM) and a nucleotide regenerating system containing acetate kinase (25 µg ml^−1^) and acetyl phosphate (15 mM), or GMPCPP (0.2 mM), were added. Reactions were centrifuged at 160,000 x *g* for 30 min in a Beckman TLA 120.1 rotor. Pellets and supernatants were resuspended in equal volumes, analyzed by SDS-PAGE and Coomassie staining, and quantified by densitometry using NIH ImageJ.

To analyze phospholipid recruitment of FtsZ polymers by FtsA, reaction mixtures were assembled containing reaction buffer, FtsZ (3 or 6 µM), GTP (2 mM) a nucleotide regenerating system containing acetate kinase (25 µg ml^−1^) and acetyl phosphate (15 mM), and, where indicated MinC (5 µM), incubated for 3 min at 23°C, and added to preassembled reactions containing FtsA or FtsA(E14R) (0 to 8 µM), SUVs (250 µg ml^−1^) and ATP (4 mM). Small unilamellar vesicles (SUVs) from *E. coli* extracts (Avanti Polar lipids) were prepared as described previously [7]. Reactions were incubated for an additional 10 min at 30°C and centrifuged at 21,130 x *g* for 15 min to pellet vesicles. Pellets and supernatants were resuspended in equal volumes, analyzed by SDS-PAGE and Coomassie staining, and quantified by densitometry using NIH ImageJ.

### Fluorescence microscopy

To observe fluorescence of FtsA in vivo, MG1655 Δ*araE*::kan, MG1655, or MG1655 *ΔminC* containing plasmid pSEB293 [4] encoding Gfp-FtsA or variants Gfp-FtsA(E14R), Gfp-FtsA(ΔMTS), or Gfp-FtsA(E14R/ΔMTS) were grown overnight on solid LB media containing ampicillin (100 µg ml^−1^), restreaked and grown for 6 hrs at 30 °C. Cells containing plasmid expressing Gfp-FtsA(ΔMTS), or Gfp-FtsA(E14R/ΔMTS) were restreaked onto LB agar plates supplemented with 0.001% L-arabinose. Cells were harvested from the plate, resuspended in PBS (phosphate buffered saline), applied to 5% agarose gel pads containing M9 minimal media supplemented with 0.2% glucose, and a coverslip was added. Samples were visualized with a Zeiss LSM 700 confocal fluorescence microscope with excitation and emission at 488/508 nm for Gfp, 515/640 nm for FM 4-64, and 358/461 nm for DAPI respectively. Where indicated, a Nomarski prism was used to acquire differential interference contrast (DIC) images. All images were captured on an AxioCam digital camera with ZEN 2012 software. Cell lengths were measured using ImageJ. The percentage of midcell fluorescence per cell was measured in ImageJ for at least 10 cells.

To observe fluorescent FtsA(ΔMTS) and FtsZ polymers, H_6_-Gfp-FtsZ and FtsZ were mixed in a 1:1 ratio for a final concentration of 8 µM in assembly buffer with GMPCPP (0.2 mM) and resuspended Talon Cobalt beads (Takara) and added, where indicated, to pre-assembled reactions of Alexa Fluor 594 labeled FtsA(ΔMTS) mixed in a 1:1 ratio with FtsA(ΔMTS) for a final concentration of 8 µM, and ATP (4 mM), where indicated. Reactions were imaged by epifluorescence microscopy using excitation and emission wavelengths of 488 and 508 nm respectively for Gfp (green channel) and 594 and 617 nm respectively for Alexa Fluor 594 (red channel) and where indicated, a Nomarski prism was used to acquire differential interference contrast (DIC) images. Lipid concentrations of FtsA and mutant proteins were measured using the lipophilic dye probe FM 4-64 as described previously [7].

### Temperature sensitive growth assays for function in vivo

To assay for temperature sensitive growth, MCA12 or MCA27 [49] containing plasmid pQE9, pQE9-*ftsA*, or pQE9-*ftsA* mutagenized to pQE9-*ftsA*(E14R), pQE9-*ftsA*(ΔMTS), or pQE9-*ftsA*(E14R/ΔMTS) were grown overnight at 30 °C in liquid LB media supplemented with ampicillin (100 µg ml^−1^), diluted to an OD_600_ of 0.1 in LB media, grown to an OD_600_ of 0.4 at 30°C, spot plated log dilutions onto LB with ampicillin agar plates, and grown overnight at 30 or 42°C, where indicated.

For immunoblotting of Gfp-FtsA, MG1655 (Δ*araE*) containing plasmid pSEB293 [4] encoding Gfp-FtsA, or Gfp-FtsA mutagenized to Gfp-FtsA(E14R), Gfp-FtsA(ΔMTS), or Gfp-FtsA(E14R/ΔMTS) was grown in liquid LB medium supplemented with ampicillin (100 µg ml^−1^) and for cells expressing Gfp-FtsA(ΔMTS), or Gfp-FtsA(E14R/ΔMTS) supplemented with 0.001% L-arabinose, to an OD_600_ of 0.8 at 30 °C. Proteins were precipitated with 15% trichloroacetic acid (Sigma-Aldrich) for 30 minutes at 4 °C. Suspensions were then centrifuged at 5,000 x *g* for 10 min at 4°C. Pellets were isolated and washed with acetone for 10 min at 4 °C followed by centrifugation at 10,000 x *g* for 10 min at 4 °C. 0.2 M NaOH was added to the pellets and then resuspended in buffer consisting of 20 mM Tris (pH 7.5) and 2% SDS, and a BCA protein determination was done. Equal concentrations of lysate (5 µg) were analyzed by reducing SDS-PAGE and transferred to a nitrocellulose membrane (Invitrogen). Membranes were washed with tris buffered saline (pH 7.6) and Tween-20 (0.05%) (TBST), blocked for 2 hours with 2% (w/v) bovine serum albumin, probed with rabbit Gfp polyclonal antibody serum and goat anti-rabbit IgG coupled with horse radish peroxidase (HRP). Gfp-FtsA and mutant proteins were visualized using Pierce ECL Western blotting substrate.

### Light scattering assays

To monitor PL remodeling dependent light scatter, reaction mixtures (80 µl) containing reaction buffer and FtsA or FtsA(E14R) (2 µM) were monitored for 5 min to collect a baseline then ATP (4 mM) or buffer was added, and reactions were monitored for an additional 60 min. To monitor FtsA(ΔMTS) polymerization dependent light scatter, reaction mixtures (80 µl) containing assembly buffer, where indicated, EDTA (15 mM) and FtsA(ΔMTS) (0 to 20 µM) were monitored at 37°C unless otherwise indicated for 5 min to collect a baseline then ATP (4 mM), ADP (4 mM) or buffer was added and reactions were monitored for an additional 30 min.

## Data availability

All data pertinent to this work are contained within this manuscript or available upon request. For requests, please contact Jodi Camberg at the University of Rhode Island.

## Acknowledgements

We thank Janet Atoyan for sequencing and microscopy assistance, Cathy Trebino, Colby Ferreira, Eric DiBiasio, Negar Rahmani and Ben Piraino for helpful suggestions and edits. Microscopy and sequencing were performed at the Rhode Island Genomics and Sequencing Center, which is supported in part by the National Science Foundation (MRI Grant No. DBI-0215393 and EPSCoR Grant No. 0554548 & EPS-1004057), the US Department of Agriculture (Grant Nos. 2002-34438-12688, 2003-34438-13111, and 2008-34438-19246), and the University of Rhode Island. The TEM data was acquired at the RI Consortium for Nanoscience and Nanotechnology, a URI College of Engineering core facility partially funded by the National Science Foundation EPSCoR, Cooperative Agreement #OIA-1655221.

## Funding

Research reported in this publication was supported in part by the National Institute of General Medical Sciences of the National Institutes of Health under Award Number R01GM118927 to J. Camberg. The content is solely the responsibility of the authors and does not necessarily represent the official views of the National Institutes of Health or the authors’ respective institutions.

## Author contributions

J.J.M. and J.C. designed and performed the experiments. J.J.M., J.C. and J.L.C. conceptualized the study and analyzed the results. J.J.M. performed electron microscopy.

J.J.M. and J.L.C. wrote the manuscript. J.L.C. obtained the funding for the study, supervised the study and managed the project. All authors reviewed, edited and approved the manuscript

## Notes

### Competing Interest Statement

The authors have declared no competing interest.

## References

[1] Haeusser DP, Margolin W. Splitsville: structural and functional insights into the dynamic bacterial Z ring. Nature reviews Microbiology. 2016;14:305–19.

[2] Liu Z, Mukherjee A, Lutkenhaus J. Recruitment of ZipA to the division site by interaction with FtsZ. Molecular microbiology. 1999;31:1853–61.

[3] Pichoff S, Lutkenhaus J. Unique and overlapping roles for ZipA and FtsA in septal ring assembly in Escherichia coli. EMBO J. 2002;21:685–93.

[4] Pichoff S, Lutkenhaus J. Tethering the Z ring to the membrane through a conserved membrane targeting sequence in FtsA. Mol Microbiol. 2005;55:1722–34.

[5] Mosyak L, Zhang Y, Glasfeld E, Haney S, Stahl M, Seehra J, et al. The bacterial cell-division protein ZipA and its interaction with an FtsZ fragment revealed by X-ray crystallography. EMBO J. 2000;19:3179–91.

[6] Hale CA, de Boer PA. Direct binding of FtsZ to ZipA, an essential component of the septal ring structure that mediates cell division in E. coli. Cell. 1997;88:175–85.

[7] Conti J, Viola MG, Camberg JL. FtsA reshapes membrane architecture and remodels the Z-ring in Escherichia coli. Molecular microbiology. 2018;107:558–76.

[8] Szwedziak P, Wang Q, Freund SM, Lowe J. FtsA forms actin-like protofilaments. The EMBO journal. 2012;31:2249–60.

[9] Loose M, Mitchison TJ. The bacterial cell division proteins FtsA and FtsZ self-organize into dynamic cytoskeletal patterns. Nature cell biology. 2014;16:38–46.

[10] Du S, Lutkenhaus J. Assembly and activation of the Escherichia coli divisome. Molecular microbiology. 2017;105:177–87.

[11] Tsang MJ, Bernhardt TG. Guiding divisome assembly and controlling its activity. Current opinion in microbiology. 2015;24:60–5.

[12] Du S, Lutkenhaus J. At the Heart of Bacterial Cytokinesis: The Z Ring. Trends in microbiology. 2019;27:781–91.

[13] Soderstrom B, Mirzadeh K, Toddo S, von Heijne G, Skoglund U, Daley DO. Coordinated disassembly of the divisome complex in Escherichia coli. Molecular microbiology. 2016.

[14] Busiek KK, Eraso JM, Wang Y, Margolin W. The early divisome protein FtsA interacts directly through its 1c subdomain with the cytoplasmic domain of the late divisome protein FtsN. Journal of bacteriology. 2012;194:1989–2000.

[15] Corbin BD, Geissler B, Sadasivam M, Margolin W. Z-ring-independent interaction between a subdomain of FtsA and late septation proteins as revealed by a polar recruitment assay. Journal of bacteriology. 2004;186:7736–44.

[16] Vega DE, Margolin W. Direct Interaction between the Two Z Ring Membrane Anchors FtsA and ZipA. Journal of bacteriology. 2019;201.

[17] Du S, Pichoff S, Lutkenhaus J. FtsEX acts on FtsA to regulate divisome assembly and activity. Proceedings of the National Academy of Sciences of the United States of America. 2016;113:E5052–61.

[18] Park KT, D. S, Lutkenhaus J. Essential Role for FtsL in Activation of Septal Peptidoglycan Synthesis. mBio. 2020;11.

[19] Dominguez R, Holmes KC. Actin structure and function. Annu Rev Biophys. 2011;40:169–86.

[20] Hanson J, Lowy J. Molecular basis of contractility in muscle. Br Med Bull. 1965;21:264–71.

[21] Kilmartin JV, Adams AE. Structural rearrangements of tubulin and actin during the cell cycle of the yeast Saccharomyces. The Journal of cell biology. 1984;98:922–33.

[22] Pollard TD, Cooper JA. Actin, a central player in cell shape and movement. Science. 2009;326:1208–12.

[23] Shaevitz JW, Gitai Z. The structure and function of bacterial actin homologs. Cold Spring Harbor perspectives in biology. 2010;2:a000364.

[24] van den Ent F, Lowe J. Crystal structure of the cell division protein FtsA from Thermotoga maritima. The EMBO journal. 2000;19:5300–7.

[25] Stoddard PR, Williams TA, Garner E, Baum B. Evolution of polymer formation within the actin superfamily. Molecular biology of the cell. 2017;28:2461–9.

[26] Cooper JA, Buhle EL, Jr., Walker SB, Tsong TY, Pollard TD. Kinetic evidence for a monomer activation step in actin polymerization. Biochemistry. 1983;22:2193–202.

[27] Gershman LC, Newman J, Selden LA, Estes JE. Bound-cation exchange affects the lag phase in actin polymerization. Biochemistry. 1984;23:2199–203.

[28] Gurung R, Yadav R, Brungardt JG, Orlova A, Egelman EH, Beck MR. Actin polymerization is stimulated by actin cross-linking protein palladin. The Biochemical journal. 2016;473:383–96.

[29] Ostrowska Z, Moraczewska J. Cofilin - a protein controlling dynamics of actin filaments. Postepy Hig Med Dosw (Online). 2017;71:339–51.

[30] Merino F, Pospich S, Funk J, Wagner T, Kullmer F, Arndt HD, et al. Structural transitions of F-actin upon ATP hydrolysis at near-atomic resolution revealed by cryo-EM. Nature structural & molecular biology. 2018;25:528–37.

[31] Chou SZ, Pollard TD. Mechanism of actin polymerization revealed by cryo-EM structures of actin filaments with three different bound nucleotides. Proceedings of the National Academy of Sciences of the United States of America. 2019;116:4265–74.

[32] Yan K, Pearce KH, Payne DJ. A conserved residue at the extreme C-terminus of FtsZ is critical for the FtsA-FtsZ interaction in Staphylococcus aureus. Biochemical and biophysical research communications. 2000;270:387–92.

[33] Pichoff S, Lutkenhaus J. Identification of a region of FtsA required for interaction with FtsZ. Molecular microbiology. 2007;64:1129–38.

[34] Geissler B, Shiomi D, Margolin W. The ftsA* gain-of-function allele of Escherichia coli and its effects on the stability and dynamics of the Z ring. Microbiology. 2007;153:814–25.

[35] Geissler B, Elraheb D, Margolin W. A gain-of-function mutation in ftsA bypasses the requirement for the essential cell division gene zipA in Escherichia coli. Proceedings of the National Academy of Sciences of the United States of America. 2003;100:4197–202.

[36] Bernard CS, Sadasivam M, Shiomi D, Margolin W. An altered FtsA can compensate for the loss of essential cell division protein FtsN in Escherichia coli. Molecular microbiology. 2007;64:1289–305.

[37] Pichoff S, Du S, Lutkenhaus J. Disruption of divisome assembly rescued by FtsN-FtsA interaction in Escherichia coli. Proceedings of the National Academy of Sciences of the United States of America. 2018;115:E6855–E62.

[38] Krupka M, Rowlett VW, Morado D, Vitrac H, Schoenemann K, Liu J, et al. Escherichia coli FtsA forms lipid-bound minirings that antagonize lateral interactions between FtsZ protofilaments. Nature communications. 2017;8:15957.

[39] Martos A, Monterroso B, Zorrilla S, Reija B, Alfonso C, Mingorance J, et al. Isolation, characterization and lipid-binding properties of the recalcitrant FtsA division protein from Escherichia coli. PloS one. 2012;7:e39829.

[40] Krupka M, Cabre EJ, Jimenez M, Rivas G, Rico AI, Vicente M. Role of the FtsA C terminus as a switch for polymerization and membrane association. mBio. 2014;5:e02221.

[41] Nag D, Chatterjee A, Chakrabarti G. FtsA-FtsZ interaction in Vibrio cholerae causes conformational change of FtsA resulting in inhibition of ATP hydrolysis and polymerization. International journal of biological macromolecules. 2020;142:18–32.

[42] Rueda S, Vicente M, Mingorance J. Concentration and assembly of the division ring proteins FtsZ, FtsA, and ZipA during the Escherichia coli cell cycle. J Bacteriol. 2003;185:3344–51.

[43] Pichoff S, Shen B, Sullivan B, Lutkenhaus J. FtsA mutants impaired for self-interaction bypass ZipA suggesting a model in which FtsA’s self-interaction competes with its ability to recruit downstream division proteins. Molecular microbiology. 2012;83:151–67.

[44] Pichoff S, Du S, Lutkenhaus J. The bypass of ZipA by overexpression of FtsN requires a previously unknown conserved FtsN motif essential for FtsA-FtsN interaction supporting a model in which FtsA monomers recruit late cell division proteins to the Z ring. Molecular microbiology. 2015;95:971–87.

[45] Du S, Pichoff S, Lutkenhaus J. Roles of ATP Hydrolysis by FtsEX and Interaction with FtsA in Regulation of Septal Peptidoglycan Synthesis and Hydrolysis. mBio. 2020;11.

[46] Bork P, Sander C, Valencia A. An ATPase domain common to prokaryotic cell cycle proteins, sugar kinases, actin, and hsp70 heat shock proteins. Proceedings of the National Academy of Sciences of the United States of America. 1992;89:7290–4.

[47] Yim L, Vandenbussche G, Mingorance J, Rueda S, Casanova M, Ruysschaert JM, et al. Role of the carboxy terminus of Escherichia coli FtsA in self-interaction and cell division. Journal of bacteriology. 2000;182:6366–73.

[48] Gayda RC, Henk MC, Leong D. C-shaped cells caused by expression of an ftsA mutation in Escherichia coli. Journal of bacteriology. 1992;174:5362–70.

[49] Dai K, Xu Y, Lutkenhaus J. Cloning and characterization of ftsN, an essential cell division gene in Escherichia coli isolated as a multicopy suppressor of ftsA12(Ts). Journal of bacteriology. 1993;175:3790–7.

[50] Fowler WE, Aebi U. A consistent picture of the actin filament related to the orientation of the actin molecule. The Journal of cell biology. 1983;97:264–9.

[51] Viola MG, LaBreck CJ, Conti J, Camberg JL. Proteolysis-Dependent Remodeling of the Tubulin Homolog FtsZ at the Division Septum in Escherichia coli. PloS one. 2017;12:e0170505.

[52] LaBreck CJ, Conti J, Viola MG, Camberg JL. MinC N- and C-Domain Interactions Modulate FtsZ Assembly, Division Site Selection, and MinD-Dependent Oscillation in Escherichia coli. Journal of bacteriology. 2019;201:e00374–18.

[53] de Boer PA, Crossley RE, Rothfield LI. A division inhibitor and a topological specificity factor coded for by the minicell locus determine proper placement of the division septum in E. coli. Cell. 1989;56:641–9.

[54] Camberg JL, Hoskins JR, Wickner S. The interplay of ClpXP with the cell division machinery in Escherichia coli. J Bacteriol. 2011;193:1911–8.

[55] Bernhardt TG, de Boer PA. SlmA, a nucleoid-associated, FtsZ binding protein required for blocking septal ring assembly over Chromosomes in E. coli. Molecular cell. 2005;18:555–64.

[56] Bisson-Filho AW, Hsu YP, Squyres GR, Kuru E, Wu F, Jukes C, et al. Treadmilling by FtsZ filaments drives peptidoglycan synthesis and bacterial cell division. Science. 2017;355:739–43.

[57] Caldas P, Lopez-Pelegrin M, Pearce DJG, Budanur NB, Brugues J, Loose M. Cooperative ordering of treadmilling filaments in cytoskeletal networks of FtsZ and its crosslinker ZapA. Nature communications. 2019;10:5744.

[58] McCullagh M, Saunders MG, Voth GA. Unraveling the mystery of ATP hydrolysis in actin filaments. J Am Chem Soc. 2014;136:13053–8.

[59] Matsui T, Han X, Yu J, Yao M, Tanaka I. Structural change in FtsZ Induced by intermolecular interactions between bound GTP and the T7 loop. The Journal of biological chemistry. 2014;289:3501–9.

[60] Pazos M, Peters K, Casanova M, Palacios P, VanNieuwenhze M, Breukink E, et al. Z-ring membrane anchors associate with cell wall synthases to initiate bacterial cell division. Nature communications. 2018;9:5090.

[61] Hill NS, Buske PJ, Shi Y, Levin PA. A moonlighting enzyme links Escherichia coli cell size with central metabolism. PLoS genetics. 2013;9:e1003663.

[62] Baker JL, Courtemanche N, Parton DL, McCullagh M, Pollard TD, Voth GA. Electrostatic interactions between the Bni1p Formin FH2 domain and actin influence actin filament nucleation. Structure. 2015;23:68–79.

[63] Busiek KK, Margolin W. A role for FtsA in SPOR-independent localization of the essential Escherichia coli cell division protein FtsN. Molecular microbiology. 2014;92:1212–26.

[64] Camberg JL, Viola MG, Rea L, Hoskins JR, Wickner S. Location of dual sites in E. coli FtsZ important for degradation by ClpXP; one at the C-terminus and one in the disordered linker. PloS one. 2014;9:e94964.

[65] Camberg JL, Hoskins JR, Wickner S. ClpXP protease degrades the cytoskeletal protein, FtsZ, and modulates FtsZ polymer dynamics. Proceedings of the National Academy of Sciences of the United States of America. 2009;106:10614–9.

[66] Conti J, Viola MG, Camberg JL. The bacterial cell division regulators MinD and MinC form polymers in the presence of nucleotide. FEBS letters. 2015;589:201–6.

[67] Blattner FR, Plunkett G, 3rd, Bloch CA, Perna NT, Burland V, Riley M, et al. The complete genome sequence of Escherichia coli K-12. Science. 1997;277:1453–62.

[68] Meyer HH, Shorter JG, Seemann J, Pappin D, Warren G. A complex of mammalian ufd1 and npl4 links the AAA-ATPase, p97, to ubiquitin and nuclear transport pathways. The EMBO journal. 2000;19:2181–92.

