## Supporting figures for "Coregulated assembly of actin-like FtsA polymers with FtsZ during Z-ring formation and division in *Escherichia coli*"

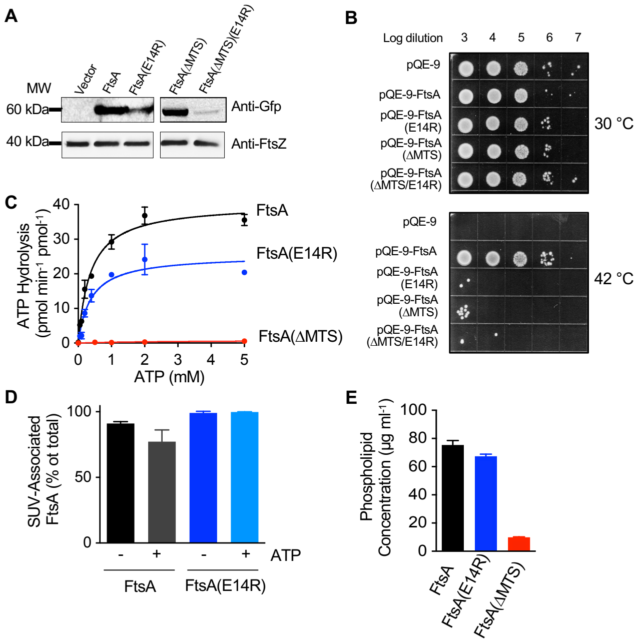


**Fig. S1. Glu 14 is important for FtsA function.** (A) Protein extracts (5 μg) from MG1655 (JC0390) cells expressing Gfp-FtsA, Gfp-FtsA(E14R), Gfp-FtsA(ΔMTS), and Gfp-FtsA(ΔMTS)(E14R) were immunoblotted with anti-Gfp and anti-FtsZ antisera as shown and described in Materials and Methods. (B) Log dilutions of *E. coli* MCA27 (*ftsA*27) cells expressing FtsA or FtsA mutant proteins from a plasmid (pQE9) and grown overnight at permissive (30 °C) and restrictive (42 °C) temperatures, where indicated, on LB agar plates containing ampicillin (100 μg $ml^{-1}$). Data shown is representative of three replicates. (C) Rate of ATP hydrolysis by FtsA wild type or mutant proteins (1 μM) measured with increasing concentrations of ATP, as described in Material and Methods. (D) Phospholipid recruitment assay with reaction mixtures of FtsA or FtsA(E14R) (2 μM), SUV's (250 μg $ml^{-1}$) and ATP then fractionated by low-speed centrifugation. (E) Purified samples of FtsA, FtsA(E14R) or FtsA(ΔMTS) (1 μM) were stained with FM 4-64 (1.25 μg $ml^{-1}$). Fluorescence was measured and compared to a standard curve of known PL concentration using SUV's. Data from (A) is representative of three replicates. Data in (C), (D) and (E) is representative of at least three replicates and represented as mean ± SEM. Pellets and supernatants for (D) were visualized by SDS-PAGE and quantified by densitometry.

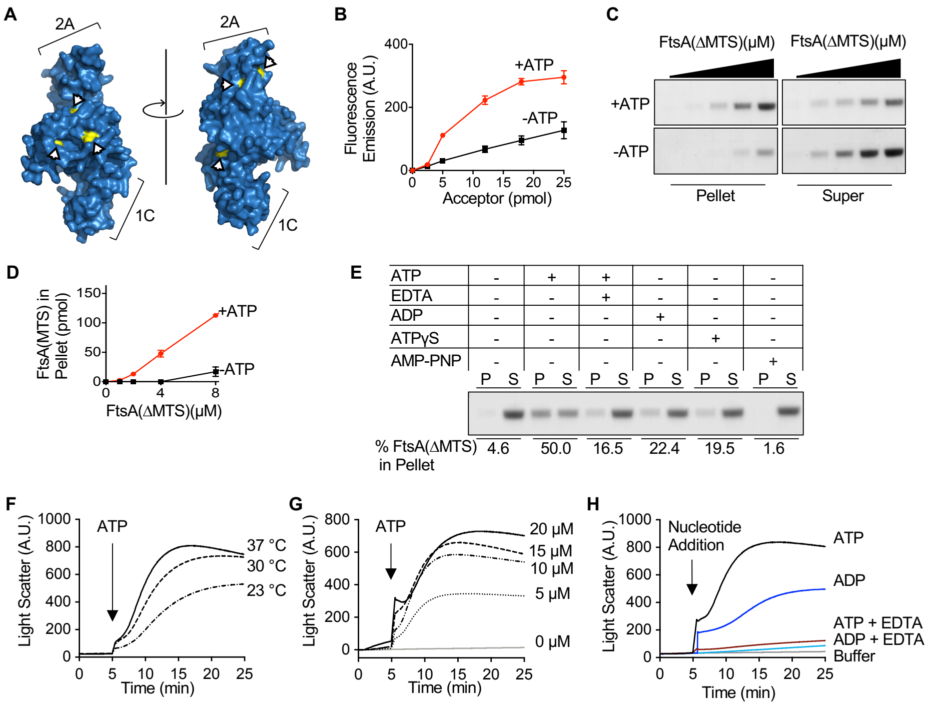


**Fig. S2. FtsA(ΔMTS) polymerization is ATP dependent.** (A) Surface representation of *E. coli* FtsA modeled onto *T. maritima* FtsA (pdb: 4A2B) [8] highlighting cysteine residues (yellow). (B) FRET using increasing concentrations of Alexa Fluor 647 labeled FtsA(ΔMTS) (2.5 to 25 pmol) were mixed with a fixed concentration of Alexa Fluor 594 labeled FtsA(ΔMTS) (100 pmol) and FtsA(ΔMTS) for a total of 600 pmol. Reactions were monitored with and without addition of ATP as described in Materials and Methods. (C) Reactions of FtsA(ΔMTS) (1, 2, 4, 8, and 12 μM) with and without ATP were incubated for 15 min and fractionated by ultracentrifugation. Pellets and supernatant were visualized by SDS-PAGE and quantified by densitometry shown in (D). (E) FtsA(ΔMTS) (8 μM) was incubated with, where indicated, ATP, ATP + EDTA, ADP, ATPγS, or AMP-PNP for 15 minutes and fractionated by ultracentrifugation. Pellets and supernatant were visualized by SDS-PAGE and quantified by densitometry and the mean of % FtsA(ΔMTS) in the pellet is shown. (F) Reactions of FtsA(ΔMTS) (15 μM) were incubated at 23, 30, and 37 ºC and monitored by 90º light scatter for 5 min to collect a baseline then ATP was added and the signal was monitored for an additional 20 min. (G) Reaction mixtures of FtsA(ΔMTS) (0 to 20 μM) were monitored as in (F) at 37 ºC. (H) Reactions of FtsA(ΔMTS) (15 μM) were monitored as in (F) at 37 ºC but after the addition of ATP, ADP, ATP + EDTA, ADP + EDTA, or buffer. Data in (B) and (D) is shown as mean ± SEM. Data from (C), (E), (F), (G), and (H) is representative of three replicates.

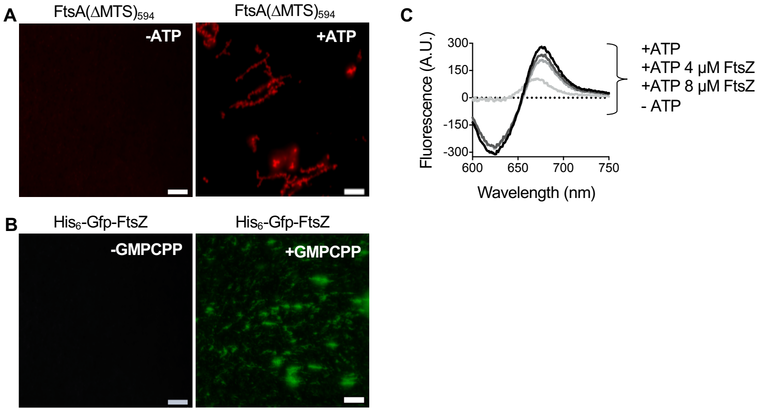


**Fig. S3. Interaction of FtsZ with FtsA(ΔMTS) polymers.** (A) Alexa Fluor 594 labeled FtsA(ΔMTS) mixed in a 1:1 ratio with FtsA(ΔMTS) and ATP for a final concentration of 8 μM. Scale bars are 5 μm. (B) $H_6$-Gfp-FtsZ and FtsZ were mixed in a 1:1 ratio for a final concentration of 8 μM with GMPCPP. Scale bars are 5 μm. (C) FRET using FtsZ (0,4 and 8 μM) mixed with Alexa Fluor 647 labeled FtsA(ΔMTS) (25 pmol), Alexa Fluor 594 labeled FtsA(ΔMTS) (100 pmol) and FtsA(ΔMTS) (475 pmol) and monitored after addition of ATP as described in Materials and Methods. Reactions in (A) and (B) were imaged by epifluorescence microscopy using excitation and emission wavelengths of 492 and 508 nm respectively for Gfp and 594 and 617 nm respectively for Alexa Fluor 594. Data from (C) is representative of three replicates.

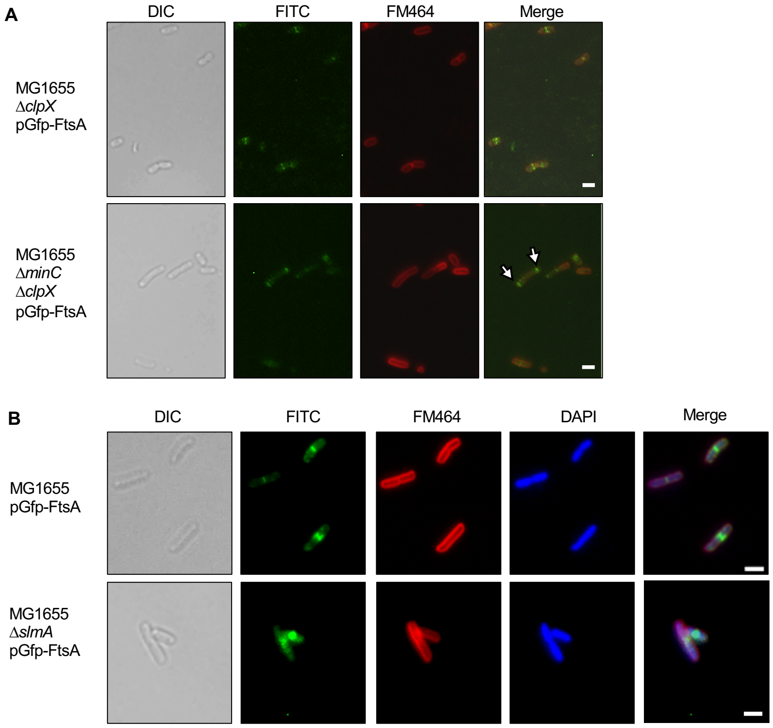


**Fig. S4. Gfp-FtsA localization in *E. coli* MG1655 Δ*clpX* and Δ*slmA* deletion strains.** Fluorescence and DIC microscopy of (A) *E. coli* MG1655 Δ*clpX,* MG1655 Δ*minC* Δ*clpX*, (B) MG1655, and MG1655 Δ*slmA* cells expressing Gfp-FtsA (green) and stained with FM-464 (red) and, where indicated, DAPI (blue). White arrows indicate polar localization. Scale bars are 2 μm.
